# A shifting role of thalamocortical connectivity in the emergence of large-scale functional brain organization during early lifespan

**DOI:** 10.1101/2024.03.11.584415

**Authors:** Shinwon Park, Koen V. Haak, Stuart Oldham, Hanbyul Cho, Kyoungseob Byeon, Bo-yong Park, Phoebe Thomson, Haitao Chen, Wei Gao, Ting Xu, Sofie Valk, Michael P. Milham, Boris Bernhardt, Adriana Di Martino, Seok-Jun Hong

**Author notes:** Corresponding author: Seok-Jun Hong.

## Abstract

While cortical patterning has been a perennial research topic in neuroscience, the mechanism for its consequence, namely functional specialization at the macro scale, remains an open question in the human brain. Here, we focused on age-dependent changes of resting-state thalamocortical connectivity to investigate its role in the emergence of large-scale functional networks across infancy, childhood and young adulthood. We found that the thalamocortical connectivity during infancy reflects an early differentiation of sensorimotor networks and genetically-influenced axonal projection. This initial role of the thalamus, however, seems to change during childhood, by establishing connectivity with the salience network and decoupling externally- and internally-oriented functional processes. Developmental simulation and perturbation analyses corroborated these findings, demonstrating the highest contribution of thalamic connectivity, especially in the later age of youth, in the formation of key characteristics of the mature brain, such as functional gradient and cortical hierarchy. Our study highlights a developmentally shifting role of the thalamus in orchestrating complex brain organization and its potential implications for developmental conditions characterized by compromised internal and external processing.

---

The human brain begins its complex developmental process soon after conception, starting as a smooth, initially uniform neural tube that bends, folds, and expands to eventually become the characteristic wrinkles of the neocortex. Over the course of pregnancy, the neural tube is patterned along the rostrocaudal axis driven by a gradient of morphogens and transcription factors ^1,2^. Simultaneously, thalamocortical axons begin to migrate into the cortical plate ^3,4^. The awareness of this intermingled process gave rise to two influential views — the Protomap and Protocortex theories — on the developmental course of how the neocortex differentiates into distinct functional regions, guided by genetic or thalamic influence, respectively ^5–7^. The current consensus is that both intrinsic (*i.e.,* genes) and extrinsic (*i.e.,* thalamus) mechanisms collectively drive cortical development, yet viewing the role of the thalamus exclusively as a mediator of sensory experience may be an oversimplification ^8–10^.

Indeed, on one hand, as most sensory information passes through the thalamus before reaching the neocortex, the thalamus is in a strategic location for mediating external signals that influence cortical differentiation ^9,10^. For instance, exposure to external sensory stimuli accelerates initial synaptic plasticity of the newborn brain and gradually shapes functional specialization through intensive interactions with the thalamus ^11–13^. On the other hand, intrinsic genetic and molecular factors also underlie the formation of thalamic nuclei as well as growth of thalamocortical projections ^14–17^. Relying on axon guidance molecules and transcription factors, thalamic nuclei topographically project to specific cortical areas, and after reaching their destination, successfully terminate their growth ^18^. While these series of processes are crucial for regional differentiation of the neocortex, exactly how the thalamocortical connectivity is involved in the neocortical functional specialization across infancy through young adulthood remains as an open question.

A core tenet of neocortical functional specialization is that it follows a sensory-association axis (SA axis), where maturation of low-level primary sensory regions precede the expansion of higher-order association areas, such as the default-mode and frontoparietal regions ^19–23^. This principle is also reflected in the development of thalamocortical connectivity, where connections with sensorimotor and salience networks are already established in neonates, while those with the frontal regions and default mode network develop later on ^24–26^. However, as the thalamus is composed of several dozen nuclei, it has been innately difficult to understand how it is globally embedded within the macroscale cortical landscape, beyond regional functional properties of individual sub-nuclei. Notably, a series of recent connectopic gradient mapping techniques based on dimensionality reduction ^27,28^ provide a viable solution, as they summarize high dimensional connectivity profiles of the brain regions (either the whole brain or specific cortical areas such as the thalamus) into parsimonious axes representing their core functional organization.

Previous studies using this technique have reported that the hierarchical structure embedded within the SA axis captures a gradient from externally-oriented processing based on perception and action on one end to stimulus-independent, internally-oriented processing on the other end ^19,29^. Interestingly, a recent study showed that this neocortical organization shows an age-dependent shift, in which the primary functional axis mainly anchored within the unimodal cortex during childhood (*i.e.,* between sensorimotor and visual regions) changes into the adult-like hierarchical gradient during adolescence ^21^. However, how the early thalamocortical interaction influences this large-scale organization across the age has not been mapped. In particular, whether thalamocortical gradients follow a similar hierarchical organization to cortico-cortical gradients (or rather show a subcortex-specific connectivity pattern across the developmental stages), and what computational principles underlie thalamus-mediated brain network generation, are largely unknown.

The current study investigated the macroscale organization of thalamocortical connectivity and its potential role in the formation of cortical functional networks during infancy and childhood/young adulthood. Continuous axes of thalamocortical connectivity were first extracted with the connectopic gradient mapping method and analyzed along the developmental age groups. Associations between cortical genes and thalamic neocortical projection maps were then characterized using gene ontology enrichment analyses. Finally, we utilized a computational approach, namely generative network modeling, to simulate the role of both thalamus and genetic information in the assembly of large-scale functional networks.

Through this multi-method approach, we found: 1) During infancy, the thalamus lays the foundation for a preliminary hierarchical network, as well as interacts with cortical genes involved in developmental processes; 2) By middle childhood and throughout adolescence/young adulthood, thalamocortical connectivity projections undertake an unique role in differentiating between internally- and externally-oriented functional processes. Indeed, the thalamic connectivity towards the salience network serves as an anchor that segregates the externally-oriented networks such as dorsal attention, visual and sensorimotor networks from internally-oriented systems such as the default mode network; 3) Generative network modeling indicated that when synthetic networks are constructed based on thalamocortical connectivity, they show the highest resemblance to networks derived from empirical data. Moreover, perturbations of key thalamo-salience network connections in different stages of development revealed the nuanced picture of underlying mechanisms of how the large-scale functional organization emerges during development. Taken together, initially there seems to be a balanced contribution of thalamus and developmentally related genes in the specialization of functional networks. However, this balance slowly changes as entering into youth, in which the thalamus steps up to a presiding position of integrating operations related to processing information from the external world and incorporating it into an internal working model.

## Results

We utilized two independent datasets across different age groups from the Human Connectome Project (HCP ^30,31^), that consists of those in infancy (n=195, 107 male; mean [SD] age=39.7 [3.04] weeks; Developing HCP, dHCP ^32,33^) and childhood/young adulthood (n=603, 417 male, mean [SD] age=14.8 [3.88] years; HCP-Development, HCP-D ^34^). Details on participant inclusion, image processing, connectopic gradient extraction, transcriptomic analysis and computational modeling are provided in **Method**, which are also schematically summarized in **Figure 1** (see the legend for a short description of our general methods).

**Figure 1.**
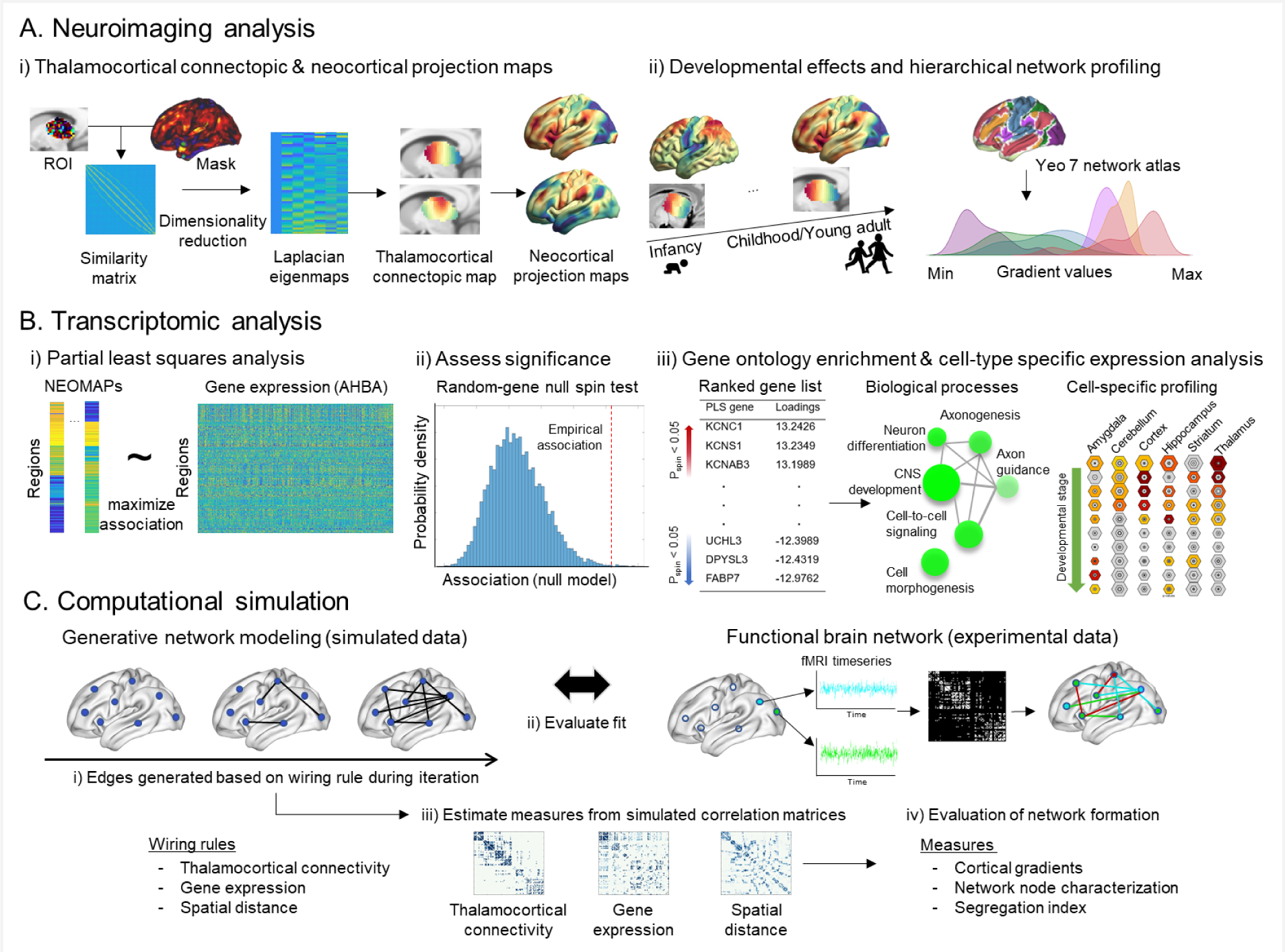
Analytical flows of (A) neuroimaging, (B) transcriptomics, and (C) computational simulation. (A-i) Connectopic mapping, a dimensional reduction technique, was used to summarize complex thalamocortical connectopic topologies. (A-ii) Aging effects were then tested for the thalamocortical maps within infancy and childhood/young adulthood groups. The maps for both the thalamus and neocortex were profiled according to the Yeo-Krienan 7 Network Atlas ^35^. (B-i) Associations between the neocortical projection maps and spatial pattern of gene expression derived from the Allen Human Brain Atlas ^36^ were quantified using partial least squares (PLS) analysis. (B-ii) Significance was tested using spatial autocorrelation-preserving permutation tests (10,000 spins). (B-iii) The significant gene sets identified by PLS analysis were ranked and then submitted to a gene ontology enrichment analysis to characterize the underlying biological processes and cell-type specific expressions across developmental stages. (C) Generative network models were utilized to investigate the effects of thalamocortical, genetic, spatial, and topological constraints when estimating how wiring costs shape functional connectome topology.

### 1. Thalamocortical connectopic mapping

To delineate the macroscale organization of thalamocortical connectivity, we extracted its connectopic gradients and neocortical projection maps using the Congrads pipeline (https://github.com/koenhaak/congrads) ^27^. In short, it applies the Laplacian Eigenmap dimensionality reduction technique to a similarity matrix of the functional connectivity between the thalamus and cortex. The resulting thalamocortical connectopic maps (CMAP) topographically represent how similar or dissimilar the thalamocortical connectivity patterns are across the thalamic voxels, while the neocortical projection maps (NEOMAPs) show a cortical counterpart to the CMAP (*i.e.,* which thalamic area is mostly connected to which neocortical area?). To ensure comparability, the individual CMAPs were aligned to the group template CMAPs using the Procrustes alignment algorithm.

#### 1.1. Thalamocortical connectopic gradients

The group-level CMAPs (“functional gradients of the thalamus”; 2 CMAPs were chosen in the study based on the variance explained; see **Supplementary Figure 1** for the details) are shown for both infancy (29-44 weeks; **Figure 2A**) and childhood/young adulthood (8-22 years; **Figure 2B**). The primary CMAP (CMAP 1) shows a gradient spanning from the anterior (red) to posterior (blue) ends of the thalamus. This pattern is relatively consistent across the two age groups (top panel of **Figure 2A, 2B**) and conveys an approximated differentiation between low-level and high-order functional networks according to the Yeo 7 network thalamic parcellation ^37^ (**Supplementary Figure 1**). The secondary CMAP (CMAP 2) comprises a gradient from superior to inferior axis during infancy (bottom panel of **Figure 2A**), and while the overall direction (superior-to-inferior) remains the same in childhood/young adulthood, the inferior part of the gradient tends to diverge into both anterior and posterior regions (bottom panel of **Figure 2B**).

**Figure 2.**
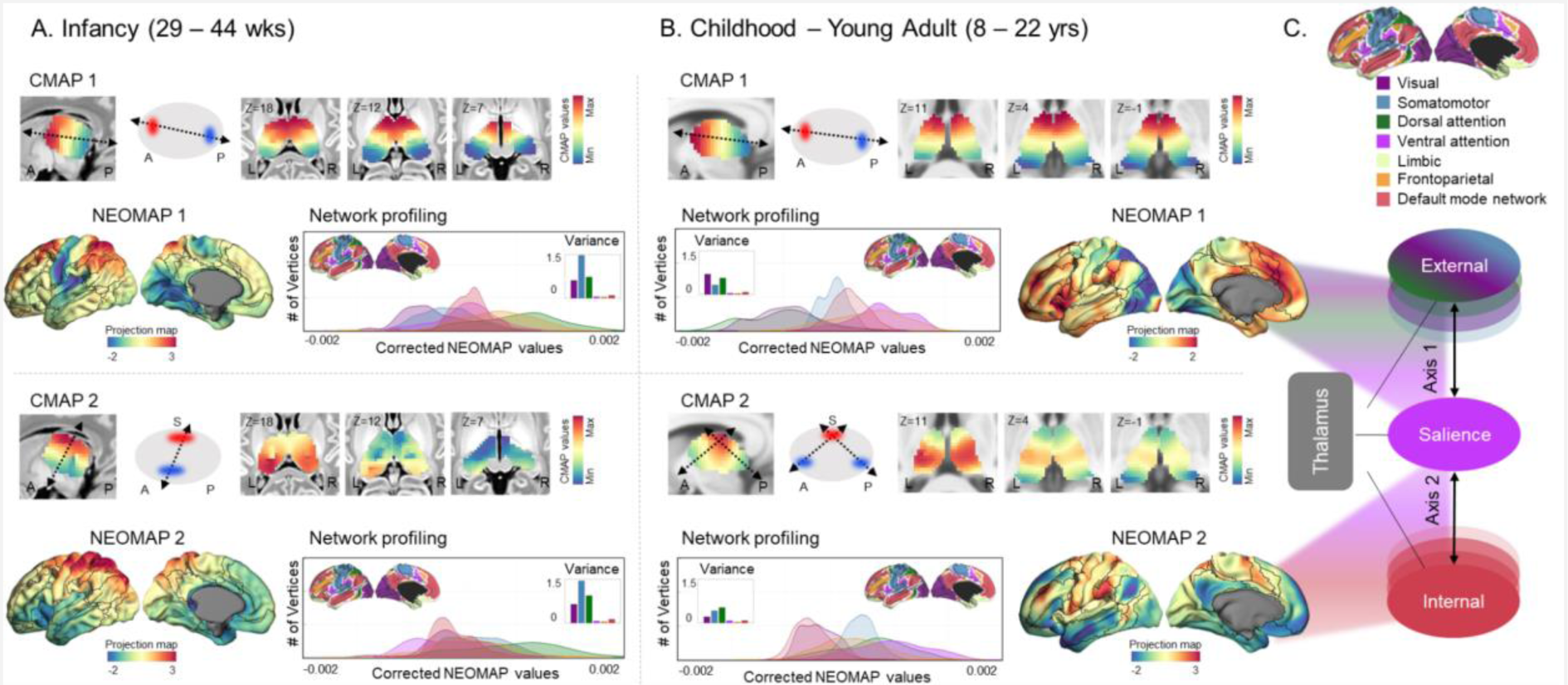
Thalamic connectopic maps (CMAP) and its neocortical projection maps (NEOMAP). CMAP 1 & 2 and NEOMAP 1 & 2 are shown for (A) infancy and (B) childhood/young adulthood. Network profiling is shown in the joy plots located next to the NEOMAPs for each panel using the Yeo-Krienan 7 Network Atlas ^35^. The bar plots embedded within each joy plot indicate the variance for each functional network. (C) A schematic of the external-to-internal axis division derived from the childhood/young adulthood NEOMAPs is depicted along with the Yeo-Krienan 7 Network Atlas legend.

It should be noted that the HCP datasets we analyzed for infancy were obtained during sleep, while those for childhood/young adulthood were collected during awake states. To investigate this potential state-dependent confounding effect in our findings, we further analyzed a separate dataset consisting of the scans taken during both sleep and awake states within individuals, of which details can be found in the **Supplementary Material**. In short, the different acquisition states (sleep vs. wakefulness) affected the magnitude of the gradient values, but not the overall topological pattern itself. In other words, the spatial patterns of the gradients were highly consistent across the acquisition conditions, which confirmed that the present findings (age-dependent different CMAPs) are not due to state differences but can be interpreted as true developmental changes. The examples of the CMAP consistency between states are presented individually (**Supplementary Figure 2**). In a similar vein, given that one of the main functions of the thalamus is regulating arousal and activity levels, we conducted the same analysis after global signal regression and found that our results were replicated (**Supplementary Figure 3**).

#### 1.2. Thalamic neocortical projection maps

The NEOMAP (*i.e.,* the neocortical counterpart of thalamus CMAP) of infancy and childhood/young adulthood showed overall diverging patterns across age (**Figure 2**), which indicates a developmental change in the relationship between the thalamus and neocortical functional organization.

In infancy, the first NEOMAP showed strong anchors in the low-level sensory and visual areas while the other end of the gradient reveals undifferentiated patterns across the brain areas (top panel of **Figure 2A**). The second NEOMAP portrayed a strong geometric pattern that seems to directly reflect the dorsoventral pattern of the second thalamic CMAP (bottom panel of **Figure 2A**). Meanwhile, in childhood/young adulthood, the NEOMAPs show relatively more differentiated patterns compared to that of infancy, gradually revealing the emergence of mature functional network systems (**Figure 2B**). This can be inferred from the network profiling results as the NEOMAP values show less variance and are more confined within each specific functional network (see joy plots and embedded bar plots in the middle of **Figure 2**). Indeed, both the first and second NEOMAPs commonly show that the salience network is situated at one end of the gradient, whereas the opposite ends of the gradients are distinct (**Figure 2B**). Specifically, the first map extends towards externally-oriented regions, such as the dorsal attention and primary sensory networks (top panel of **Figure 2B**), whereas the second map mainly is occupied by the internally-oriented default mode network (bottom panel of **Figure 2B**). Taken together, the thalamocortical projections appear to contribute to a large-scale organization in which the salience network serves as a stable anchor for differentiation of external and internal axes (**Figure 2C**).

#### 1.3. Thalamic effects in segregation of the external-to-internal axis

To further confirm the above hypothesis, we quantified two different network-level indices reflecting – i) internal-external axis segregation and ii) an association with core/matrix thalamic organization. First, we created a segregation index by contrasting the salience network with both externally and internally oriented functional areas, as this network is recognized for its function to switch between various large-scale networks ^38^. Specifically, *Segregation_SAL-EXT_* indicates the difference of averaged NEOMAP between the salience network and networks comprising externally oriented functional processes including the dorsal attention and visual networks. Meanwhile, *Segregation_SAL-INT_* quantifies the degree of isolation between the salience and default mode network. Changes in the *Segregation* indices were measured along the 5 different age groups with equally spaced age bins as follows: infancy (< 37 weeks (wks), n=49; 37-40 wks, n=59; 40 wks, n=47; 41 wks, n=48; 42-44 wks, n=52); childhood/young adulthood (8-10 years (yrs), n=81; 10-13 yrs, n=125; 13-16 yrs, n=173; 16-20 yrs, n=141; 20-22 yrs, n=83) (**Figure 3A**). As expected, there was greater segregation between the salience network and both externally- and internally-oriented networks during childhood/young adulthood in comparison to infancy. This suggests that the external-internal axis more prominently emerges during childhood/young adulthood (**Figure 3A**).

**Figure 3.**
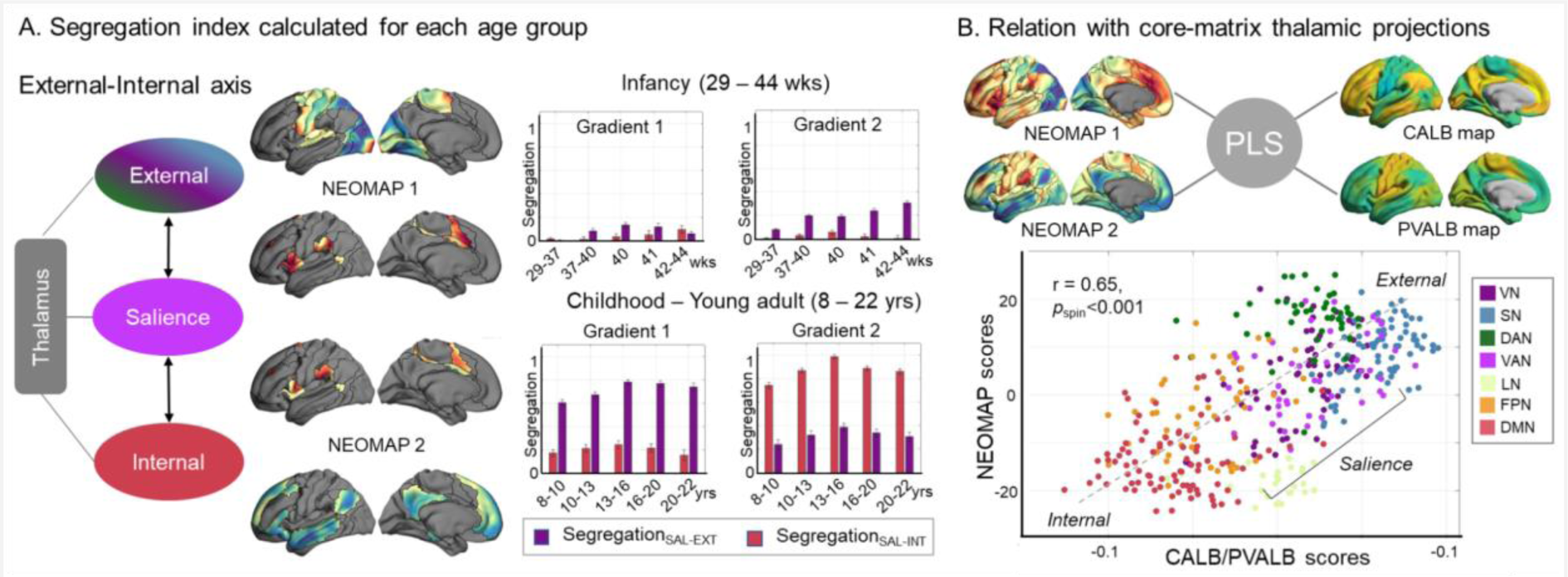
Thalamocortical influence on cortical network segregation. (A) The external-internal segregation is quantified and profiled along age subgroups. The differentiation between external-internal axis does not clearly emerge until childhood/young adulthood periods. The first NEOMAP represents segregation of the salience network with externally-oriented networks while the second NEOMAP shows segregation with the internally-oriented networks. (B) Association between childhood/young adulthood NEOMAPs and core-matrix thalamic projections. Abbreviation. NEOMAP, Neocortical projection map; SAL, salience; EXT, external; INT, internal; PLS, Partial least squares method, CALB, Calbindin; PVALB, Parvalbumin

Compared to externally-oriented networks that tend to be more regionally confined (*e.g.,* visual and somatosensory networks), those for internal processes seem to recruit widely distributed areas across the cortex ^39^. This differentiation may reflect distinct patterns of underlying thalamic projections based on the relative density of ‘core’ and ‘matrix’ cells ^10,40^. Specifically, on the one hand, Parvalbumin (PVALB)-rich core cells are typically found in sensory thalamic nuclei and send axonal projections to concentrated areas of primary sensory cortices. On the other hand, calbindin (CALB)-rich matrix cells are more abundant in higher-order thalamic nuclei and project on the cortex in a more dispersed manner.

Corroborating this previous finding, the partial least squares analyses also found significant association between the NEOMAPs and cytoarchitecture-based thalamic projections of core and matrix cells. Moreover, correlation between the latent variables reflects the external-to-internal axis with the salience network situated in the middle of the distribution (**Figure 3B**).

#### 1.4. Age-dependent changes in CMAPs & NEOMAPs

The thalamocortical CMAPs and NEOMAPs per se also showed significant aging effects (**Supplementary Figure 4**). Specifically, in infancy, at each end of the CMAPs, an overall expansion (*i.e.,* more functional differentiation) seems to occur with age. A similar pattern is also depicted in the NEOMAPs. In childhood/young adulthood, fairly consistent aging effects are found as well, although the slope was lower compared to that of infancy. Contrary to the infancy group which showed mainly a gradient expansion, the childhood/young adulthood period presented both expansion and convergence with age. A supplementary video showing an average change in gradient values across age is included (**Supplementary Video**).

### 2. Transcriptomic analyses of thalamocortical projection maps

By far, we have shown that the thalamocortical connectivity makes widespread neocortical projections with distinct patterns reflecting a large-scale functional organization of cortical networks. In brief, the first infancy NEOMAP revealed a preliminary differentiation between low-level sensory and high-order transmodal regions, whereas the NEOMAPs of childhood/young adulthood collectively depicted a segregation between functional cortical networks involved in externally-vs internally-oriented processes. Compared to these, however, the second NEOMAP of infancy (bottom panel of **Figure 2A**) was not clear on its developmental implication, although its graded expression along the dorsoventral axis resembles the characteristic areal patterning of the developing neocortex driven by morphogens and transcription factors. Thus, we examined its association with cortical gene expression, based on Allen Brain Human Atlas (ABHA) and the partial-least squares (PLS) analysis, an unsupervised multivariate method. To contextualize the multivariate patterns with respect to biological processes, we conducted a subsequent gene ontology analysis with *ShinyGo 0.77* (http://bioinformatics.sdstate.edu/go) ^41^. Finally, we did a cell-type specific expression analysis (CSEA) (http://genetics.wustl.edu/jdlab/csea-tool-2) ^42^, which uses the BrainSpan dataset (http://www.brainspan.org) to compare developmental expression profiles across multiple age spans.

The associations between gene expression and the second infancy NEOMAP revealed a single latent variable that was significant, after a spin test that preserves spatial autocorrelation (*r*=0.64; *P*_spin_=0.02).The latent variable explained 41.4% of the NEOMAP variance. The post hoc loading analysis revealed a total of 850 strongly contributing genes with both negative and positive loadings that showed significant contribution to the identified latent variable. Further gene enrichment analysis revealed that the candidate negative and positive genes were largely involved in different underlying biological processes. Importantly, processes involved in the development of the central nervous system, ranging from axon/neuron projection guidance and neurogenesis to various morphogenesis-involved differentiation, were mainly driven by the negative down-regulated genes (**Figure 4A**, **Supplementary Figure 5, Supplementary Table 1**). These results were replicated using another gene ontology enrichment analysis tool – *Enrichr* ^43–45^ (Supplementary Figure 6). Further comparison of developmental expression profiles of these genes across specific cell-types using CSEA showed concentrated expression in the thalamus and cortex, from early fetal throughout infancy (**Figure 4B**). The positive up-regulated genes belonged to pathways of cellular component disassembly and various transmembrane transport processes (**Supplementary Table 2**). Finally, for completion, we tested genetic associations for the other NEOMAPs, and either found no significant enrichment or if there was, those genes were not associated with developmental processes (**Supplementary Figure 7**).

**Figure 4.**
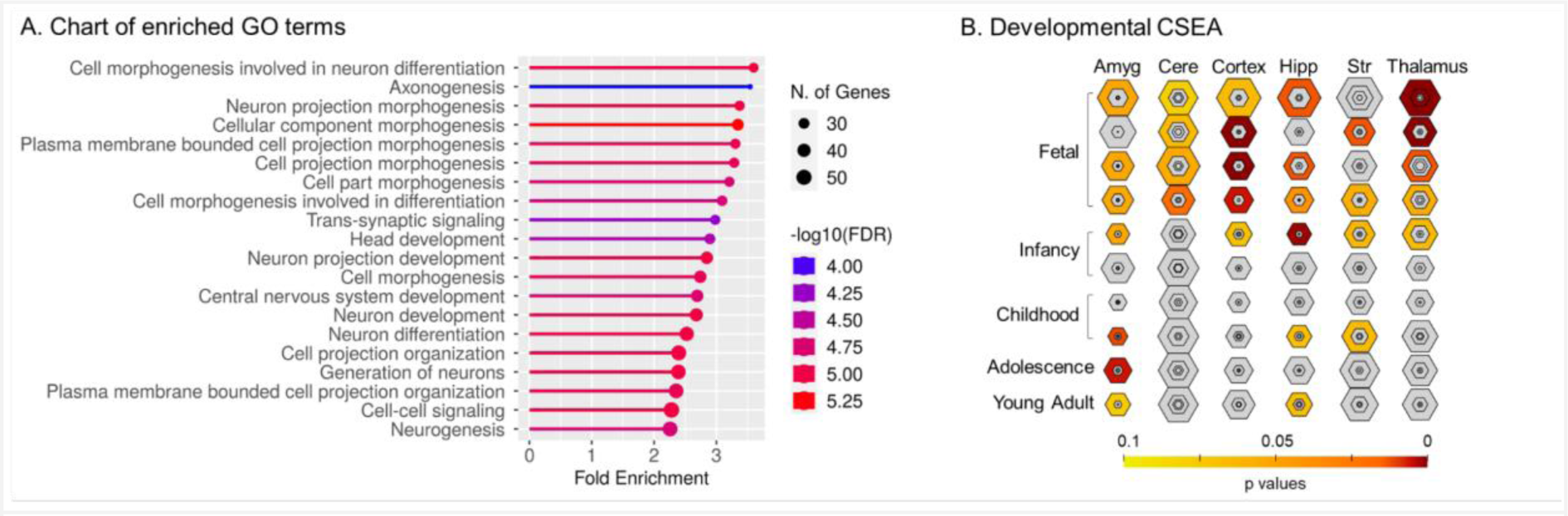
Top enriched gene ontology (GO) terms indicating biological processes of genes showing significant association with the negatively enriched genes of the second NEOMAP in infancy. (A) Fold enrichment is a measure of the degree to which a particular term is overrepresented in the genes of interest compared to a background set of genes. FDR indicates how likely the enrichment represents a false positive finding. (B) For the cell-type specific expression analysis (CSEA), genes enriched in a particular cell type were identified by specificity index p-value (pSI) across four thresholds (0.05, 0.01, 0.001, 0.0001), where lower pSI values indicate more specific genes. The size of the hexagons from outside to center correspond to the pSI thresholds (0.05, 0.01, 0.001, 0.0001), respectively. Colors indicate the FDR-adjusted P-values.

### 3. Computational modeling analysis

Finally, to gain a mechanistic understanding of whether and how thalamocortical connectivity contributes to the growth of cortical networks, we further explored a series of ‘network generative models’, which creates synthetic networks according to predefined wiring rules to mimic developmental processes ^46^. Our main interests in this analysis were to compare the relative effects of the thalamocortical connectivity and gene expression in functional network generation. For comparison of model performance, we also tested the effects of relevant spatial and topological features, as per previous studies ^47,48^. Model performance was assessed based on the Kolmogorov-Smirnov (KS) statistic that quantified the difference between synthetic and empirical networks based on node degree, node clustering, node betweenness, and edge length distribution ^47^. Detailed explanation for each of the wiring rules (*i.e.,* thalamocortical connectivity, gene expression, spatial, and topological factors) can be found in the Methods section.

In general, the models based on developmental factors (*i.e.,* gene expression and thalamocortical connectivity) performed better compared to the spatial and topological factors (**Figure 5A, Supplementary Figure 8**). The thalamocortical connectivity-based rules particularly yielded the best fitting to the empirical data of Childhood/Young adults (Childhood/Young adult: mean *KS*_thal_=0.09, *KS*_gene_=0.16, *KS*_spatial_=0.45; *p*_corrected_<0.0003; **Figure 5A**). Contrarily, gene expression and thalamocortical connectivity rules showed similar performance in Infancy (Infancy: *KS*_thal_=0.20; *KS*_gene_=0.19; *KS*_spatial_=0.43; **Figure 5A**). After comparing model performance, we further reconstructed the adjacency matrices derived from the simulations and then used the mean of individual adjacency matrices to extract cortical gradients and calculate network modularity as well as the segregation index (*i.e.,* within – between network functional connectivity). Notably, the gradients derived from the synthetic networks built according to the thalamocortical connectivity rule generated remarkably similar patterns of the canonical cortical gradients during childhood/young adulthood (**Figure 5B**). Also, significantly higher network modularity was found in the synthetic networks based on thalamocortical connectivity compared to the other two networks during childhood/young adulthood (*Q*=0.53, *p*_corrected_<0.0003; **Figure 5C**). Finally, corroborating our experimental finding (in section 1.3), the thalamocortical model showed the largest segregation index between the salience and externally-oriented networks (*i.e.,* visual, somatosensory, dorsal attention networks) as well as the internally-oriented default mode network during childhood/young adulthood (**Figure 5D**).

**Figure 5.**
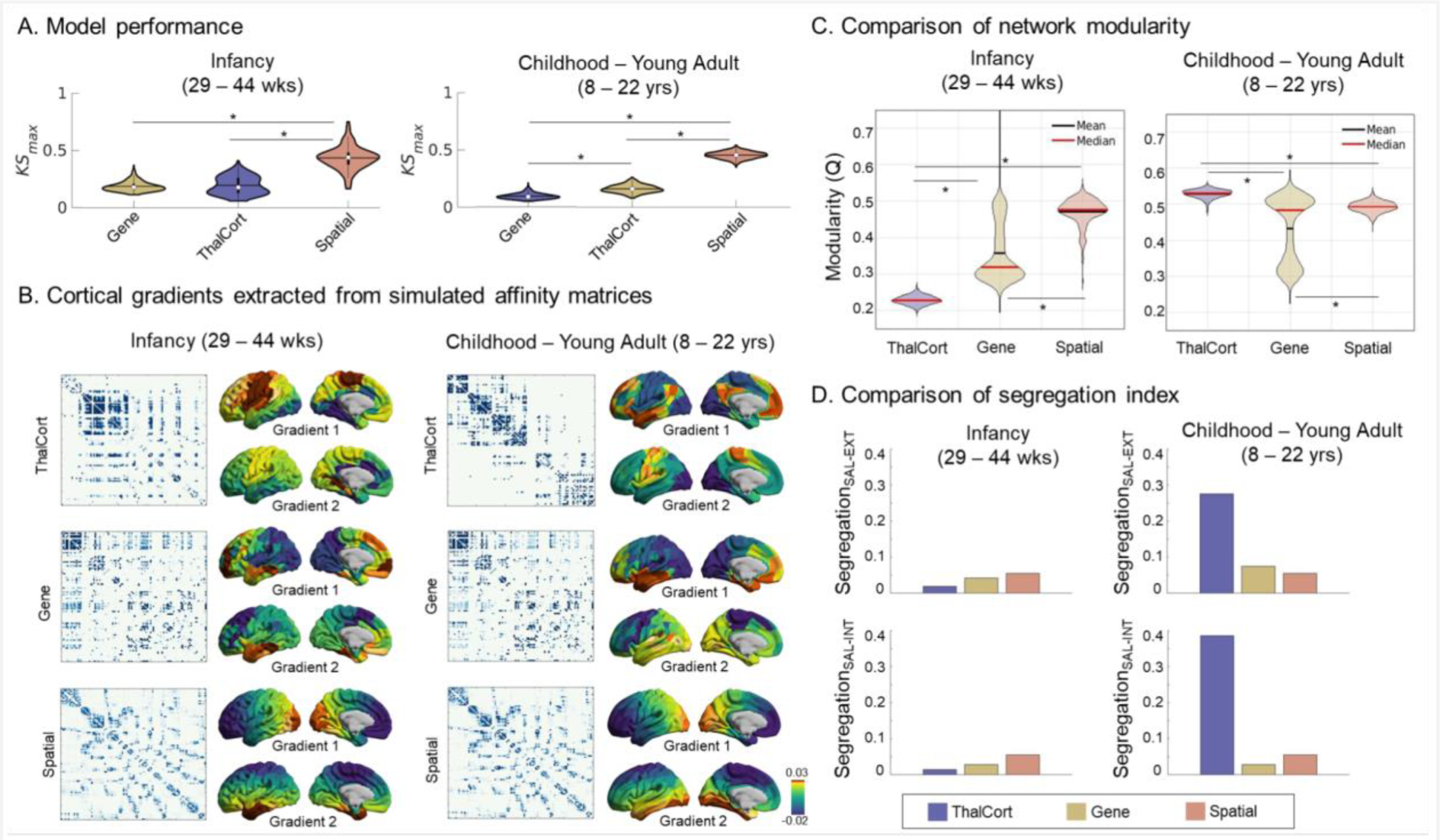
Generative network modeling. (A) Model performance is measured using Kolmogorov-Smirnov statistics, where lower values imply greater similarity to empirical data. (B) The first two cortical gradients extracted from the affinity matrices of the synthetic networks are shown for each different wiring rule. Comparison of (C) network modularity and (D) segregation indices calculated from each synthetic network based on different wiring rules. * indicates a significant pairwise difference at p-values adjusted for Bonferroni correction (*p*<0.0003). Abbreviation: Thalcort, Thalamocortical connectivity; Gene, correlated gene expression; SAL, salience network; EXT, externally-oriented networks; INT, internally-oriented networks; w, weeks; y, years

In sum, we have substantiated the importance of thalamocortical connectivity in the emergence of large scale organization – segregation between external and internal processes. Next, to further enhance a mechanistic understanding of how thalamocortical connectivity affects the functional organization across different ages, we performed perturbation to thalamo-salience connectivity in a series of growth models (an approach informing the network simulation with age-dependent developmental information; see **Figure 6A** and **Method** for the details). In comparison to a no-perturbation growth model, a total of 4 perturbation models were tested: 1) perturbation applied to the 8-12 yrs age group; 2) 12-16 yrs age group; 3) 16-22 yrs age group; 4) all age groups. As expected, perturbations compromised the segregation of both internal and external axes, but notably, the former (*i.e.,* the salience-internal) was more dramatically affected, suggesting a higher contribution of thalamocortical connectivity on the development of internal processing areas (default mode network) during youth (**Figure 6B**). In particular, the perturbation had a profound impact at a later stage of development (>12yrs), when most higher-level cognitive functions such as abstract thinking and reasoning are actively developed ^49^. Finally, the simulated cortical gradients further corroborate the underlying mechanism of thalamo-salience connectivity and how it differs according to when it was perturbed during different developmental stages (**Figure 6C**).

**Figure 6.**
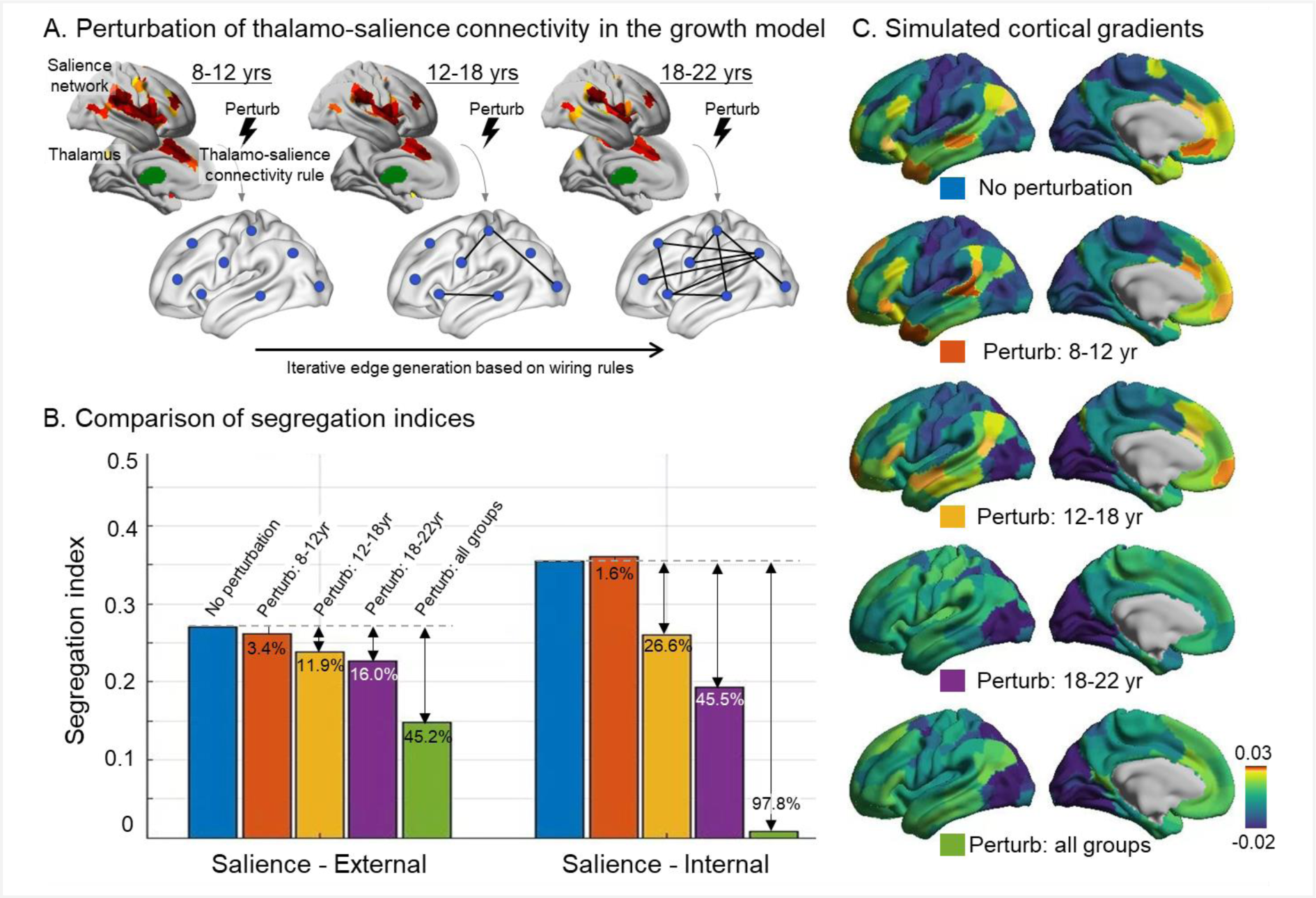
Perturbation of developmentally informed growth models. (A) A schematic of the growth model based on thalamo-salience connectivity wiring rules that changed over the age span. In comparison to a *no-perturbation* model, 4 models that perturbed the thalamo-salience connectivity in differing developmental age groups were tested as follows: 1) perturbation applied to the 8-12 yrs age group; 2) 12-16 yrs age group; 3) 16-22 yrs age group; 4) all age groups. (B) The segregation indices (salience - external; salience - internal) were calculated for each growth model. The percentage indicates the amount of difference compared to the *no-perturbation* model. (C) Cortical gradients extracted from the simulated affinity matrix of each growth model are shown.

## Discussion

Functional specialization of a large-scale brain network shows extensive changes in connectivity profiles over development before stabilizing in adulthood. In the current study, we demonstrated, both experimentally and theoretically, the importance of the thalamus in this process, revealing its age-dependent role change in associating multiple functional subsystems. Major sensory information passes through the thalamus before reaching the neocortex, naturally highlighting its role in shaping functional network organization ^50^. However, this role seems to change with ongoing maturation. Indeed, during infancy, it provides scaffolding for the marked transformation to hierarchically differentiate between low-level and higher-order regions, while also interacting with cortical genes involved in developmental processes. However, once cortical functional arrangement stabilizes into distinct networks, we found that the thalamus further engages in a system-level segregation between externally- and internally-oriented systems. These results have been also substantiated through computational simulations of generative networks. Specifically, the thalamocortical connectivity constraints not only produced the most realistic cortical networks, but also contributed to the segregation between systems and yielded even connectopic topologies that closely resemble the cortico-cortical gradients ^29^. Furthermore, perturbations to the thalamo-salience connectivity in different developmental stages had distinct effects for the external-to-internal segregation and simulated cortical gradients. Taken together, these results provide an expanding perspective for the role of thalamus in the manifestation of large-scale functional networks that we observe in the developing brain.

Previous work has established the potential effect of thalamus in the early developmental trajectory of structural organization of the brain ^16^. Indeed, microstructural properties such as myelin formation and cortical thickness develop earliest in cortical regions that receive first-order thalamic relays ^51^. Meanwhile, protracted maturation in regions of the transmodal cortex, which are heavily associated with high-order thalamic relays, are more vulnerable to prematurity and show experience-dependent plasticity ^19,52,53^. Here, we extend prior findings and reveal that the thalamus in the infant brain sends projections to the neocortex primarily anchored in sensorimotor and visual areas while those related to presumably higher-order regions are still not fully specialized ^24,54^. This delay in development may be because the formation of higher-order functional networks premises an integration of distributed regions across the cortex, which in turn requires the establishment of long-range connections and efficient synaptic pruning that occurs throughout childhood ^55–57^. Such protracted maturation of high-order systems may provide advantages by facilitating the development of primary sensory regions required for early survival and adaptation, as well as fostering experience-dependent plasticity following birth ^58–60^. In addition, our findings are in line with the established pattern of specialized connections between thalamic nuclei and cortical regions. The first NEOMAP of infants shows that low-level sensory cortical regions have high connectivity with the posterior/lateral thalamic regions, such as the pulvinar, medial geniculate nucleus, and ventroposterolateral nucleus, which are known to project to visual, auditory, and somatosensory regions, respectively ^9,61^. On the other hand, the anterior regions of the thalamus (e.g., anterior, mediodorsal, and ventral anterior nuclei), which have been linked to higher-order functions ^9^, demonstrate strong connections with the prefrontal cortex and posterior association regions.

Shortly after birth, there is an upsurge of strengthened connectivity between neurons, subsequently followed by a more prolonged selective pruning process that continues on until young adulthood ^62,63^. In this initial stage, various localized cellular and molecular mechanisms such as synaptogenesis, axonogenesis, and dendritic arborization underlie the dynamic changes pertaining to postnatal surface expansion ^64–68^. In our study, the second NEOMAP in the infant group showed associations with genes enriched for related processes such as axonogenesis, neuron differentiation as well as cell projection morphogenesis and signaling. This is in line with previous studies that support the critical role of thalamic axonal projections in the maturational process of neocortical connectivity ^69,70^. Alternatively, the tilted inferior-superior axis depicted in this projection map reflects the thalamocortical connectopic gradient pattern, which can be also interpreted in light of the Dual Origin theory. This theory conceptualizes the sequential progression of cortical differentiation as originating from the piriform cortex (paleocortex) and the hippocampus (archicortex) ^71–73^ and expanding to the sensorimotor and prefrontal cortical areas, which results similarly in the inferior-superior axis that we found from the second NEOMAP ^72,73^.

A core aspect of cortical organization that underlies cognitive development is captured in the organizing axis from sensory (*i.e.,* external) to association (*i.e.,* internal) functional areas. This principle encompasses a wide range of developmental features related to the cortical organization of cyto-architecture, structural connectivity and functional connectivity ^74^. Our study found that this fundamental principle is summarized in the “two main axes” of thalamic cortical function: i) The first NEOMAP range from salience networks to other cortical networks involved in externally-oriented processes such as the dorsal attention and low-level sensory regions. The dorsal attention and sensory networks are externally driven and phasic in nature, implying a selective attentional process to specific sensory input ^75–77^. ii) In the second NEOMAP, an internally-oriented system (*e.g.,* the default mode network), is situated at the other end of the salience networks. This gradient may represent how the thalamus contributes to shaping internal mental model building, or generative models of the environment. In fact, the retrosplenial cortex, a region closely associated with the default mode network, shows strong connections with the anterior thalamic nuclei and hippocampus, which are regions that may coordinate to update existing mental representations ^78–80^.

Of utmost importance, the salience network, a key component of both the first and second NEOMAPs, shows integrative connectivity across wider distances and diverse regions of the cortex, and thus plays a critical role in facilitating communication between multi-level systems at the global level ^81^. It has been suggested to be largely responsible for selecting the most homeostatically relevant information among the complex sensory inputs from the external world to continuously update our internal model ^82^. This process entails the coordination of brain network dynamics between externally and internally oriented cognitive processing, which the salience network is most capable of given its position in the middle of the ‘sensory-fugal’ axis ^81^. In sum, thalamocortical projections encompassing these networks may engage in the selection and gating of external inputs of high salience as a means of allocating attentional resources and updating prior beliefs.

Building upon the neuroimaging findings, we further sought to confirm whether the thalamocortical connectivity is indeed a core mechanism for a large-scale brain organization (rather than an epiphenomenon), by applying a generative network modeling approach. In previous studies, the synthetically generated networks have been shown to faithfully recapitulate major characteristics of network organization that are observed in the empirical data ^47,48,83^. In our study as well, when informing the network generation with thalamocortical connectopic patterns, the functional segregation of cortical networks naturally emerges, and its patterns were strikingly similar with those from the real brain, which was not the case in the models informed by different wiring rules. Specifically, compared to the models based on spatial distance, topological features or genetic correlations, the formation of a hierarchical brain network with meaningful functional segregation was driven only by the thalamocortical connectivity model. Notably, this pertains to modeling results from childhood/young adulthood and not infancy, which confirms our observation of diverging CMAP patterns between the two age groups. Using a growth model that allows for the tracking of brain network patterns across the iterations (analogous to developmental phases in the real brain) ^47^, we found that the thalamic influence on functional segregation is exceedingly greater during later developmental stages. Particularly, compared to the externally-oriented functional regions, segregation of the internally-oriented area is strongly modulated by thalamo-salience connectivity, implying its role in the maturation of the default mode network. Our results therefore provide a mechanistic clue of how thalamocortical connectivity affects the whole-brain connectogenesis.

Several limitations should be noted. First, our results demonstrated the underlying network configuration of thalamocortical connectivity, assuming it was static during the entire scan. However, in recent years, dynamic, non-stationary aspects of brain activity have received increasing attention, and several methods, including co-activation pattern and quasi-periodic pattern analyses ^84,85^, have been proposed for investigating these dynamic fluctuations. While previous studies have found that large-scale temporal coordination of connectivity patterns depend on highly connected hubs in the cortex, whether subcortical regions such as the thalamus play a role in cortical synchronization may be addressed in future studies. Secondly, the confounding effect of different acquisition conditions (i.e., sleep vs. awake states) limits our ability to interpret the results as being solely related to developmental effects. However, we demonstrated that the underlying architecture per se remains unchanged across different arousal states with an independent dataset of individuals who were scanned in both sleep and awake states. Future data collection efforts should consider including data acquisition in both sleep and awake states within the same neurotypical individual to enable test/retest reliability of the results across different arousal states. Finally, despite the extensive coverage of developmental periods from infancy through childhood and young adulthood (thanks to the efforts of open science and data sharing in the neuroimaging field), accessible datasets did not include adequate data for early childhood periods (ages 1-7 yrs). However, the Healthy Brain and Child Development initiative (https://heal.nih.gov/research/infants-and-children/healthy-brain) is now addressing these missing age gaps with the initiation of longitudinal, large-scale cohorts from infancy to early childhood. Such large-scale, open, collaborative efforts greatly enhance our understanding of the developing human brain and provide unprecedented opportunities for reproducibility of science.

All together, we revealed age-dependent changes in thalamocortical connectopic gradients and their corresponding neocortical projection maps. Neuroimaging and transcriptomic analyses gave us a novel insight into its shifting influences across infancy and childhood/young adulthood in macroscale functional organization, whereas computational models confirmed the critical role of thalamocortical connectivity in functional network segregation. While our study focused exclusively on the interactions between the thalamus and cortex, further studies may collectively consider the extensive connections that thalamus has with other subcortical regions such as the basal ganglia and cerebellum ^86–88^. Finally, as the thalamus is a critical region for regulating cortical networks, alterations in thalamocortical connectivity may be a harbinger of aberrant whole-brain connectivity that underlies various developmental conditions, including schizophrenia and psychosis, epilepsy, and autism ^89–94^.

## Online methods

### 1. Data acquisition and preprocessing

#### 1.1 Infancy

For infancy, we analyzed neuroimaging data from the second release of the Developing Human Connectome Project (dHCP) neonatal data (http://www.developingconnectome.org/second-data-release/). Subjects were included if both structural and functional data were available. All anatomical scans were reviewed by a neuroradiologist who provided a radiology score. Scans with a score of 3 (incidental findings with unlikely clinical significance but possible analysis significance) or 5 (incidental findings with possible/likely significance for both clinical and imaging analysis) were excluded from further analyses. After quality control and preprocessing, the final sample consisted of 195 healthy term infants (107 male, mean [SD] gestational age at birth=39.9 [1.26] weeks, mean [SD] age at scan=41.0 [1.67] weeks) and 60 pre-term infants (42 male, mean [SD] gestational age at birth=32.6 [3.12] weeks, mean [SD] age at scan=35.5 [2.78] weeks).

MR images were acquired on a 3T Philips Achieva scanner equipped with a dedicated neonatal imaging system and a neonatal 32 channel phased array head coil at the Evelina Neonatal Imaging Centre, St Thomas’ Hospital, London. All infants were scanned without sedation during natural sleep. High-resolution T2w and T1w structural images were acquired with a 0.8 mm isotropic spatial resolution and a thickness of 1.6mm, overlapped by 0.8 mm (T2w: TE/TR=156 ms/12 s; T1w: TE/TR/TI=8.7/4795/1740 ms). Resting-state fMRI data were acquired using multiband accelerated echo-planar imaging for approximately 15 minutes (TE/TR=38ms/392ms, flip angle=34°, spatial resolution=2.15 mm isotropic, 2300 volumes). Single-band reference scans were also collected with bandwidth matched readout and additional spin-echo EPI acquisitions with both AP/PA phase-encoding directions.

Preprocessed structural and functional data using the dHCP minimal preprocessing pipelines were included in the second release (for details see ^32,33^). Briefly, for structural preprocessing, motion-corrected, reconstructed T2w images were first bias-corrected and then brain extracted (using BET; ^95^) for the segmentation procedure using the Draw-EM algorithm (https://github.com/MIRTK/DrawEM, ^96^). Then, cortical white, pial, and midthickness surfaces were extracted and then inflated and projected to a sphere using a Multi-Dimensional Scaling procedure. Cortical surfaces were aligned to the 40-week dHCP surface template using Multimodal Surface Matching with higher-order constraints (^97,98^; https://github.com/ecr05/MSM_HOCR). This procedure provides one-to-one point correspondence between vertices across all subjects, which ensures that each vertex index represents the same anatomical location on multiple surfaces. Minimal preprocessing of the resting-state fMRI data included susceptibility field distortion correction, slice timing and motion correction, registration to both T2w individual structural image and 40-week group template, temporal high-pass filtering (150s high-pass cutoff), and independent component analysis (ICA) denoising using FSL-FIX. The first 10 volumes were discarded in subsequent analyses to achieve steady-state magnetization of the fMRI signal, resulting in 2290 volumes. We excluded participants with mean framewise displacement (FD) > 0.4 mm, which enables a trade-off between sample size and data quality.

#### 1.2 Childhood/Adolescence

Neuroimaging data of typically developing participants aged 8-21 years were downloaded from the repository of Lifespan HCP-Development (Release 2.0) at the National Institute of Mental Health Data Archive (NDA) ^34^. After quality control and preprocessing, the final sample consisted of 603 healthy participants (417 male, mean [SD] age=14.8 [3.88] years). Participants were excluded for insufficient English, premature birth, MRI contraindications, or a lifetime history of serious medical conditions, endocrine disorders, head injury, psychiatric/neurodevelopmental disorders.

Participants are scanned on Siemens 3T Prisma (whole-body scanner with 80mT/m gradient coil, slew rate of 200T/m/s; Siemens 32-channel head coil; software version: E11C) using matched protocols across 4 sites (Harvard University, University of California – Los Angeles, University of Minnesota, and Washington University in St. Louis) in the United States. T1-weighted images were acquired using a 3D multi-echo MPRAGE sequence (TR=2500, TI=1000 ms, TE=1.8/3.6/5.4/7.2 ms, flip angle=8°, FOV=256×240×166 mm, 320×300 matrix, 208 slices, 0.8 mm isotropic voxels). Resting-state fMRI data were acquired with a multiband gradient-recalled (GRE) EPI sequence (TR=800 ms, TE=37 ms, flip angle=52°, FOV=208 mm, 104×90 matrix, 72 oblique axial slices, 2 mm isotropic voxels, multiband acceleration factor of 8, 1912 volumes). A total of 4 fMRI scans were acquired in an eyes open with passive crosshair viewing condition for approximately 26 minutes in both the AP/PA phase-encoding directions.

We used the minimally preprocessed structural and functional data using the public release HCP pipelines v3.22 (https://github.com/Washington-University/HCPpipelines) that were released as a part of the Lifespan HCP-Development Release 2.0. Details of the specific preprocessing procedures can be found elsewhere ^30,31^. Briefly, for structural data, the HCP PreFreeSurfer pipeline performs bias-field and gradient distortion correction, alignment between T1w and T2w images, and registration to MNI space. The FreeSurfer pipeline goes through the typical FreeSurfer procedures including brain extraction, intensity normalization, followed by cortical parcellation and subcortical segmentation. Finally, the PostFreeSurfer pipeline converts the data into NIFTI and GIFTI formats that can be used as Connectome Workbench files in surface space. For functional data, the HCP fMRIVolume pipeline includes correction for gradient nonlinearity, motion, fieldmap-based EPI distortion, boundary based registration of EPI to T1w, non-linear (FNIRT) registration to MNI152 space, and intensity normalization. Following this procedure is the HCP fMRISurface pipeline, which projects the volumetric data into standard grayordinate space (which includes cortical surface vertices and subcortical voxels) and then applies smoothing with a 2mm FWHM gaussian kernel. The first 10 volumes were discarded, resulting in 1902 volumes. As with the dHCP data, FD was calculated and participants with a mean FD that exceeded 0.4 mm were excluded from the final analyses.

### 2. Data analysis

#### 2.1. Connectopic gradient analysis

We estimated thalamocortical connectopic maps (CMAP) which topographically represent dominant modes of functional connectivity change within the thalamus, based on the voxel-wise connectivity between each thalamic voxel and the rest of the neocortex. This procedure is described in detail elsewhere ^27^. In brief, we can extract the CMAP in largely two steps that consist of 1) constructing a similarity matrix followed by a 2) dimensionality reduction process. First, we prepared two time-by-voxel matrices using the fMRI time series data, each from the thalamus and the neocortex. Before computing the correlation between these two matrices, we performed a lossless dimensionality reduction in the latter due to its relatively large size, using singular value decomposition. Then we used the *η2* coefficient to quantify the similarity among the voxel-wise correlations between thalamic voxels and neocortex vertices. For the second step, we followed previous studies to apply the Laplacian eigenmaps to the resulting similarity matrix, which yielded the CMAPs within the thalamus that represent the dominant modes of change in functional connectivity between the thalamus and neocortex ^27,99^. We initially extracted the CMAPs at the group level for each dataset (dHCP, HCPD) by using the group-averaged similarity matrix, to use as a template for subsequently aligning the individual CMAPs using Procrustes alignment algorithm. Neocortical projection maps (NEOMAP) were calculated using the aligned CMAPs of each individual (https://github.com/koenhaak/congrads; lines 260 - 280 in conmap.py). In order to translate the CMAP patterns within the thalamus onto the neocortex we first obtained the similarity between the CMAP and thalamic activation pattern for each TR (=beta). Next, using these betas we computed the dot product with the neocortex timeseries activation to yield an average of the neocortical fMRI maps across the TRs, weighted by the betas. In short, the NEOMAPs were ‘color-coded’ according to that of the thalamic voxel showing the highest correlation (*e.g.,* red vertices in the NEOMAP are most highly connected with the red voxels in the CMAP) ^100^.

CMAPs and NEOMAPs were statistically tested for significant aging effects, controlling for sex and mean FD, using surface-based linear models implemented in a Matlab toolbox, SurfStat (http://www.math.mcgill.ca/keith/surfstat/; https://brainstat.readthedocs.io/en/master/) ^101,102^. Regions that showed significant positive and negative aging effects at FDR-corrected p<0.05 were delineated with black and white lines, respectively (**Supplementary Figure 3**). To interpret these gradients, we profiled the CMAP and NEOMAPs using the Yeo-Krienan 7 Network Atlas ^35,37^. Finally we divided the participants into 5 age groups for each dataset to visualize the change in gradient values across age. We divided the samples by following the age ranges representative of developmental stages reported in previous literature ^103–105^, as well as considering the distribution of participants for each group. The range and number of participants for the 5 age groups are as follows: infancy ( < 37 weeks (wks), n=49; 37-40 wks, n=59; 40 wks, n=47; 41 wks, n=48; 42-44 wks, n=52); childhood/young adulthood (8-10 years (yrs), n=81; 10-13 yrs, n=125; 13-16 yrs, n=173; 16-20 yrs, n=141; 20-22 yrs, n=83). To measure the impact of thalamus on segregation between functional networks we devised a segregation index as follows: *Segregation_SAL-EXT_* indicates the difference between NEOMAP values of the salience network and networks comprising externally oriented functional processes such as the dorsal attention, visual and somatosensory networks. On the other hand, *Segregation_SAL-INT_* quantifies the degree of isolation between the salience and default mode networks.

#### 2.2. Genetic analysis

##### 2.2.1. Microarray Gene Expression

Regional microarray expression data were obtained from 6 post-mortem brains (1 female, ages 24.0 - 57.0, mean [SD]=42.50 [13.38]) provided by the Allen Human Brain Atlas (AHBA, https://human.brain-map.org; ^36^). Data was processed with the *abagen* toolbox (version 0.1.3; https://github.com/rmarkello/abagen) using the Schaefer 400-region surface-based atlas in MNI space. However, as only two of the brains in the AHBA sampled both hemispheres, we focused our analysis on the left cortex.

First, microarray probes were reannotated using data provided by ^106^; probes not matched to a valid Entrez ID were discarded. Next, probes were filtered based on their expression intensity relative to background noise ^107^, such that probes with intensity less than the background in >=50.00% of samples across donors were discarded , yielding 31,569 probes. When multiple probes indexed the expression of the same gene, we selected and used the probe with the most consistent pattern of regional variation across donors (i.e., differential stability; ^108^), calculated with:

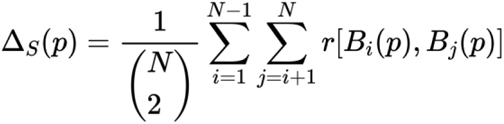

where *r* is Spearman’s rank correlation of the expression of a single probe, *p*, across regions in two donors *B_i_* and *B_j_*, and *N* is the total number of donors. Here, regions correspond to the structural designations provided in the ontology from the AHBA.

The MNI coordinates of tissue samples were updated to those generated via non-linear registration using the Advanced Normalization Tools (ANTs; https://github.com/chrisfilo/alleninf). Samples were assigned to brain regions by minimizing the Euclidean distance between the MNI coordinates of each sample and the nearest surface vertex. Samples where the Euclidean distance to the nearest vertex was more than 2 standard deviations above the mean distance for all samples belonging to that donor were excluded. To reduce the potential for misassignment, sample-to-region matching was constrained by hemisphere and gross structural divisions (*i.e.*, cortex, subcortex/brainstem, and cerebellum, such that *e.g.,* a sample in the left cortex could only be assigned to an atlas parcel in the left cortex; ^106^). If a brain region was not assigned a sample from any donor based on the above procedure, the tissue sample closest to the centroid of that parcel was identified independently for each donor. The average of these samples was taken across donors, weighted by the distance between the parcel centroid and the sample, to obtain an estimate of the parcellated expression values for the missing region. This procedure was performed for 9 regions that were not assigned tissue samples. All tissue samples not assigned to a brain region in the provided atlas were discarded.

Inter-subject variation was addressed by normalizing tissue sample expression values across genes using a robust sigmoid function ^109^:

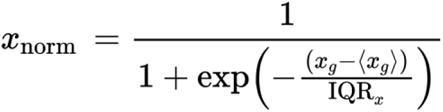

where 〈*xg*〉 is the median and IQR is the normalized interquartile range of the expression of a single tissue sample across genes. Normalized expression values were then rescaled to the unit interval:

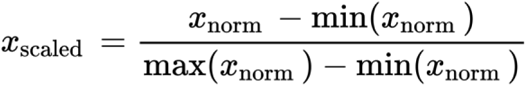

Gene expression values were then normalized across tissue samples using an identical procedure. Samples assigned to the same brain region were averaged separately for each donor and then across donors, yielding a regional expression matrix with 200 rows (=half of Schaefer 400 ROIs), corresponding to brain regions, and 15,631 columns, corresponding to the retained genes.

##### 2.2.2. PLS Analysis

We used partial least squares (PLS) analysis to investigate the correlation between the NEOMAPs and spatial pattern of gene expression derived from AHBA. PLS is an unsupervised multivariate statistical technique used to identify orthogonal sets of latent variables that maximize the covariance between the two datasets (in our case, gene expression values and NEOMAPs). The general underlying model of PLS is as follows:

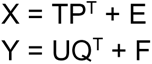

where X is a 200×15631 AHBA regional gene expression matrix and Y is a 200-element vector representing the neocortical projection map of interest. T and U are, respectively, the latent scores that are derived from the projections of X (200×1) and projections of Y (200×1), while P (15631×1) and Q (1×1) are the eigenvectors indicating the loading weights for each latent score. PLS finds the P and Q that maximize the covariance of U and T. The latent variables are obtained by applying singular value decomposition on the data after it has been z-scored.

To test whether the observed relationship between gene expression and NEOMAPs revealed by the PLS model was statistically significant, we used spatial autocorrelation-preserving permutation tests, or spin tests. The surrogate maps of spatially permuting parcellated brain regions while conserving for spatial autocorrelation were generated using the BrainSMASH python toolbox (https://brainsmash.readthedocs.io/en/latest/) ^110^. We compared the empirical correlation result between latent scores from real data to the null distributions of 10,000 correlations derived from applying PLS to the spatially permuted datasets. The same test procedure was used to test for the significance of feature contribution of each gene to the association between gene expression and NEOMAPs. Specifically, the empirical loading value of each gene was statistically tested using the null distribution 10,000 loadings obtained for each gene. Genes with positive loadings indicate up-regulation whereas the negative loadings represent a down-regulation in relation to the neocortical projection map. Genes with both negative and positive weights that showed a significance of *p_spin_*<0.05 were considered in the enrichment analysis.

Finally, by implementing a targeted gene approach, we examined whether the NEOMAPs reflect distinct patterns of thalamic projections based on the relative density of ‘core’ and ‘matrix’ cells. Each sub-nucleus of the thalamus consists of a combination of excitatory neurons with distinct cell types that project differentially to the neocortex. On the one hand, granular-projecting ‘core’ cells are prevalent in sensory thalamic nuclei and project to specific regions of the cortex such as low-level primary sensory cortices. On the other hand, supragranular-projecting ‘matrix’ cells, which are typically dense in higher-order thalamic nuclei, project more diffusively across the cortex. Using the data provided by the AHBA, we conducted a PLS analysis between the NEOMAPs and genetic expression of calcium-binding proteins, parvalbumin and calbindin, which represent core and matrix cells, respectively.

##### 2.2.3. Gene Enrichment Analyses

To characterize the biological processes of the significant gene sets identified by PLS analysis, we used ShinyGO 0.76.2 ^41^, a toolbox for testing the gene enrichment against Gene Ontology Biological Processes (Available online: http://bioinformatics.sdstate.edu/go/). The 15,631 genes retained from preprocessing the AHBA data were used as the background reference set. We explored the biological processes with which the genes are significantly involved, separately, based on their loading signs. Significant results are reported after Benjamin-Hochberg FDR correction (q<0.05). Finally, we fed the significant gene list into the cell-type specific expression analysis (CSEA) developmental expression tool (http://genetics.wustl.edu/jdlab/csea-tool-2) ^42^, which utilizes the BrainSpan dataset (http://www.brainspan.org) for constructing developmental expression profiles. The statistical significance of overlap between our candidate genes and cell-specific genes for each developmental stage was tested with Fisher’s exact test with Benjamini-Hochberg (BH) multiple testing correction for the number of cell types assayed. Genes enriched in a particular cell type were identified by specificity index p-value (pSI) across four thresholds (0.05, 0.01, 0.001, 0.0001), where lower pSI values indicate more specific genes. However, it should be noted that the significant genes were extracted from the AHBA which consists of adult postmortem data, thus, the results shown with CSEA do not directly reflect developmental processes, although many genes have been found to be consistent across the lifespan and conserved during evolution ^111,112^. Of the 550 genes, 476 genes existed in the brain region and development expression dataset.

#### 2.3. Computational modeling analysis

Finally, we sought to confirm whether our neuroimaging findings (*e.g.,* functional effects of thalamocortical connectivity) do not merely describe phenomenological aspects of the developing brain but truly reflect an underlying mechanism. To this end, we employed an advanced simulation approach, called ‘generative network modeling’. Using this tool, we were able to investigate the effects of thalamocortical, genetic, spatial, and topological constraints by synthetically creating functional connectome topology. We used the cost-topology trade-off model, which has been widely implemented to study human brain networks ^47,48,83,113–115^ and adopted the analysis code made available by Oldham, et al. (https://github.com/StuartJO/GenerativeNetworkModel). We used the Schaefer 200 cortical parcellation and focused only on the left cerebral hemisphere, to match our genetic analysis, and also to reduce computational costs due to numerous model iterations.

A total of 7 trade-off models derived from a combination of the thalamocortical, genetic, spatial, and topological constraints as parameters were tested as follows: 1) spatial model; 2) spatial + thalamocortical connectivity model; 3) thalamocortical connectivity model; 4) spatial+correlated gene expression model; 5) correlated gene expression model; 6) thalamocortical connectivity + correlated gene expression model; and 7) a model based on a topological ‘matching’ rule (**Figure 5; Supplementary Figure 8**). For the thalamocortical connectivity constraint, we used the similarity matrix of the correlation between thalamus and cortex, derived from the connectopic gradient analysis. To compute the correlated gene expression constraint, we calculated the Pearson correlation between 1,899 brain-specific genes ^116,117^, using the AHBA transcriptomic data. While there is data available for the developing brain (BrainSpan atlas; http://www.brainspan.org), the spatial coverage is inadequate to fully estimate the correlated gene expression across the whole brain. Oldham, et al. estimated both uncorrected, raw correlated gene expression as well as those corrected for their intrinsic spatial autocorrelation, and found that the former shows better performance, which motivated us to only use the uncorrected estimates of correlated gene expression ^47^. For the spatial parameter, we calculated the physical distance of a connection between brain regions using the Euclidean distance algorithm, based on the assumption that longer physical distances will require more metabolic resources to form a connection, thus reflecting wiring cost ^118^. The spatial factor imposes a distance-based constraint that forms a connection between nodes of shorter rather than longer distances. As per previous studies ^47,48^, synthetic networks formed based on this rule did not show high resemblance to the empirical network because of the lack of central hub brain regions that facilitate connections between nodes of long-distance. Thus, following previous studies, we also included a topological factor that forms a connection between nodes that share a certain topological feature ^47,48^. The matching index was included as the representative topological factor because it showed the best performance amongst other topological factors in previous studies ^47,48^. This rule estimates the similarity between a node’s neighborhood (i.e., whether it is connected to similar nodes) and forms a connection based on this overlap ^47,48^.

We evaluated the model performance by comparing the synthetic networks derived from the models to empirically extracted networks in terms of degree, clustering, betweenness, and edge length distributions. For each measure, the distributions were compared using the Kolmogorov-Smirnov (KS) statistic, a non-parametric test that quantifies the distance between two distributions, where lower values imply greater similarity. For conservative evaluation, we determined the performance of a given model based on the maximum KS statistic (corresponding to the least accuracy). Next, a 3-step multi-resolution optimization procedure that sampled 10,000 combinations of different parameters was used to obtain the best-fitting parameters for each model ^48^. To start, a random sample of 2000 points in the parameter space was selected, where each point is defined by a specific combination of the model parameters. Using these randomly sampled parameters, 2000 synthetic networks were made and evaluated using the max (KS) fit statistic. After network evaluation, a Voronoi tessellation was performed to identify regions within the parameter space that showed better fit. Then, a further 2000 points were sampled following a probability that controls which cells the points are sampled from. This step was repeated 5 times with the probability of sampling from cells with better fits increased at each repetition, yielding a total of 10,000 points to evaluate. The synthetic networks for each individual were constructed using the parameters that were chosen based on the evaluation.

Next, we wanted to estimate how well the synthetic networks made from thalamocortical connectivity models reflect the functional segregation between cortical networks as seen in the neuroimaging data analyses. The affinity matrices derived from the synthetic networks showing best performance were analyzed in terms of modularity network measure, as well as for constructing cortical gradients. The Brain Connectivity Toolbox ^119^ and the Brainspace toolbox ^28^ were used for these analyses, respectively. Notably, to avoid circularity in the evaluation of thalmic contribution to cortical network formation, measures such as cortical gradients and segregation index, which were not used for parameter estimation were employed. Finally, we tested a series of growth models based on perturbations to thalamo-salience connectivity in different developmental age groups (Childhood: 8-12 yrs, Adolescence: 12-18 yrs, Young adulthood: 18-22 yrs) to examine the underlying mechanism of how thalamocortical connectivity affects the functional organization across development. In comparison to a no-perturbation growth model, a total of 4 perturbation models were tested: 1) perturbation applied to the 8-12 yrs age group; 2) 12-16 yrs age group; 3) 16-22 yrs age group; 4) all age groups. Contrary to the previous analyses that compared the overall model performance between different biological, spatial, and topological constraints, here, we used the growth models (https://github.com/StuartJO/GenerativeNetworkModel) to provide age-dependent developmental information of the thalamocortical connectivity during network simulation. Perturbations were made to the thalamo-salience connectivity by first identifying the nodes within the salience network using the Yeo-Krienan 7 network atlas, and then randomly mixing its connections to preserve the overall amount of connectivity despite the disarranged connectivity patterns.

## Supplementary Materials

**Supplementary Figure 1.**
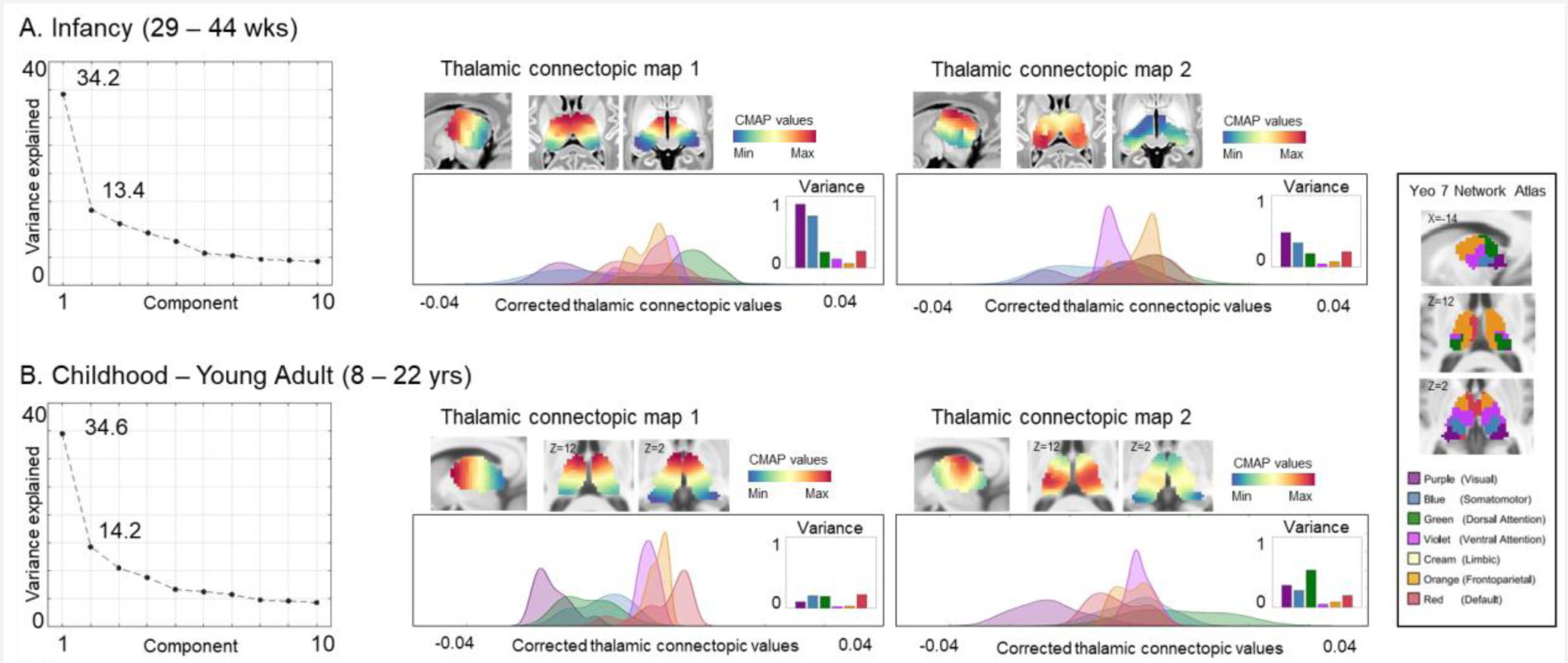
Plots showing variance explained and hierarchical network profiling of the thalamic connectopic maps using the Yeo 7 network thalamic parcellation ^37^ for (A) infancy and (B) childhood – young adulthood (parcellation from: https://github.com/ryraut/thalamic-parcellation)

### Comparison of sleep-wake states

The HCP datasets for infancy were obtained during sleep, while those for childhood/young adulthood were collected during awake states. Obtaining sleep data during infancy is a common practice to reduce motion artifacts. However, a direct comparison between these two datasets engenders a potential interpretational confound in regards to whether the results can be attributed to developmental effects, or the effects of arousal states. While some previous studies highlight the consideration of state-dependent differences in functional connectivity ^121–123^, others report that similar spatial resting-state functional connectivity maps are obtained during sleep and awake states ^124,125^.

Using a separate, within-subject dataset consisting of two MRI sessions – one in wakefulness and one in natural sleep – we investigated this potential state-dependent effect in our findings. As we could not find an open-source dataset that contains a within-subject data acquired in both sleep and wake states, we used an in-house dataset collected by one of our collaborators (A.D.M), which is currently for the purpose of another study targeting children with autism. Specifically, we analyzed resting-state fMRI scans collected from 29 children with autism (23 males) aged 5-8 years (inter-session mean time: 12±12 days), on an Allegra 3T scanner (TR=2000ms, TE=15ms, voxel size=3×3×3mm). Data with median framewise displacement (FD)≤0.2 mm were preprocessed using the Configurable Pipeline for the Analysis of Connectomes (CPAC) version 1.8.4 (https://fcp-indi.github.io/docs/latest/user/index) ^126^. After extracting the thalamic gradients, we first tested the spatial correlation between different states at the group level as well as for each individual. Then, using a mixed effects model (covariates: age, sex, medianFD), we tested for differences in gradient magnitude.

We found strikingly high correlations between the thalamic gradients extracted from sleep and awake scans on the group level (1^st^ gradient: r=0.9860; 2^nd^ gradient: r=0.9747; Supplementary Figure 2A), as well as the individual level (mean [sd] = 1^st^ gradient: 0.9411 [0.04]; 2^nd^ gradient: 0.9047 [0.04]; all individuals showed r>0.75; **Supplementary Figure 2B**). Despite the overall similarity in gradient patterns there were significant differences in gradient magnitude between sleep and awake scans (**Supplementary Figure 2C**). This implies that although there are indeed substantial state-dependent effects in the brain network formation, the effects are likely due to a different strength of network manifestation, not a complete reorganization of the essential network configuration. Therefore, the current findings based on the observation of the thalamic connectome topology (not magnitude) should be interpreted in light of developmental changes rather than attributed as differences in data acquisition (sleep *vs*. awake states).

**Supplementary Figure 2.**
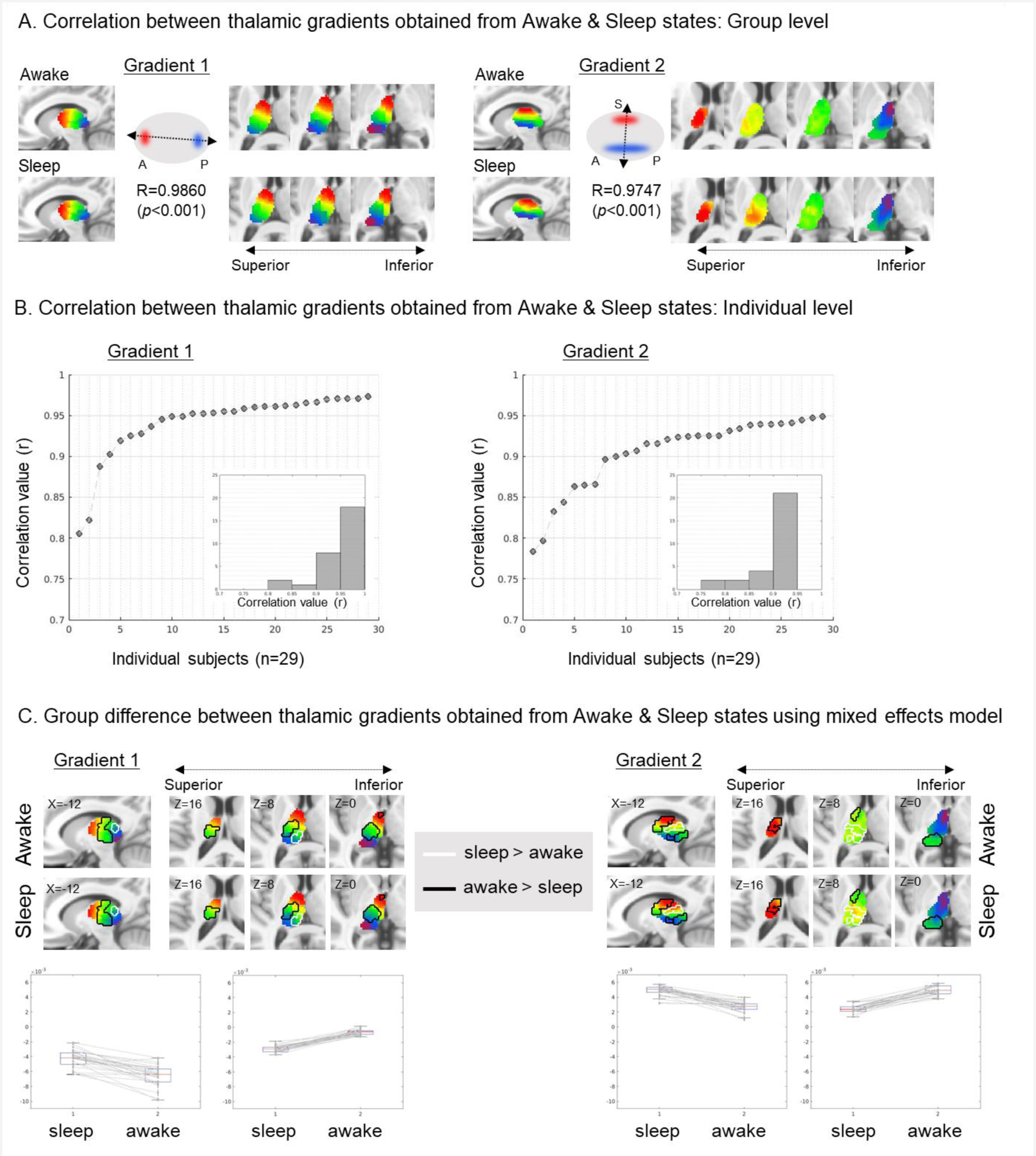
Associations between thalamic gradients obtained from awake vs sleep states collected in a separate sample. (A) Thalamic gradients are visualized at the group level for both awake and sleep states. Strong correlations are found for both gradient 1 (R=0.9860, p<0.001) and gradient 2 (r=0.9747, p<0.001). (B) Correlation values between the awake and sleep state thalamic gradients for each individual are sorted from the lowest to highest value. Majority of the individuals show a correlation higher than 0.9 for both gradients.

**Supplementary Figure 3.**
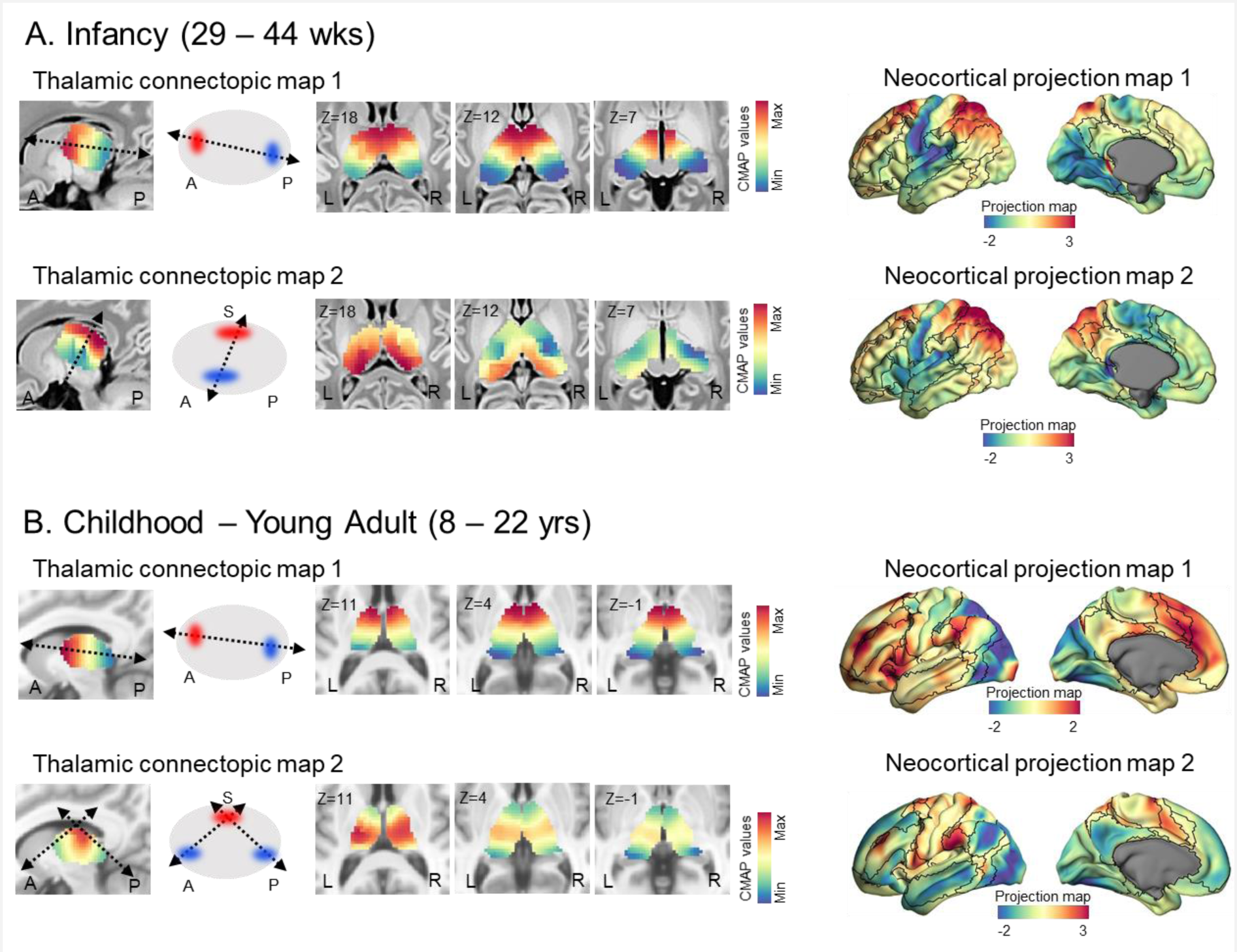
Thalamic connectopic maps (CMAP) and its projection to the neocortex (NEOMAP) after global signal regression for (A) infancy and (B) childhood/young adulthood.

### Age-related differences in thalamocortical connectopic gradient and projection maps

Beyond the simple group-level visual inspection between infant and childhood/young adulthood, we further examined the age-dependent change of both CMAPs and NEOMAPs within each group. The thalamocortical CMAPs showed significant aging effects indicating that the thalamus plays a different role in shaping neocortical functional networks across development (**Supplementary Figure 4)**. In infancy, the regions with positive gradient values (red), situated at one end of the gradient, show a significant positive aging effect, whereas areas with negative gradient values (blue) at the other end, reveal a negative aging effect (**Supplementary Figure 4A**, left; P_FDR_<0.05). In short, at each end of the gradient, an overall expansion (*i.e.,* more functional differentiation) seems to occur with age. A similar pattern is also depicted in the NEOMAPs (**Supplementary Figure 4B**, left). Again, this is also interpreted as a more distinct segregation between the low-level sensory regions and brain regions that involve high-order networks undergoing the process of differentiation. In childhood/young adulthood, fairly consistent aging effects are found, although the effect size was weaker compared to that of infancy, as seen in the scatter plots (**Supplementary Figure 4A**, right). Contrary to the infancy group which showed mainly a gradient expansion, the childhood/young adulthood period presented both expansion and convergence along the age (**Supplementary Figure 4B**, right). A supplementary video showing an average change in gradient values across age is included (**Supplementary Video**).

**Supplementary Figure 4.**
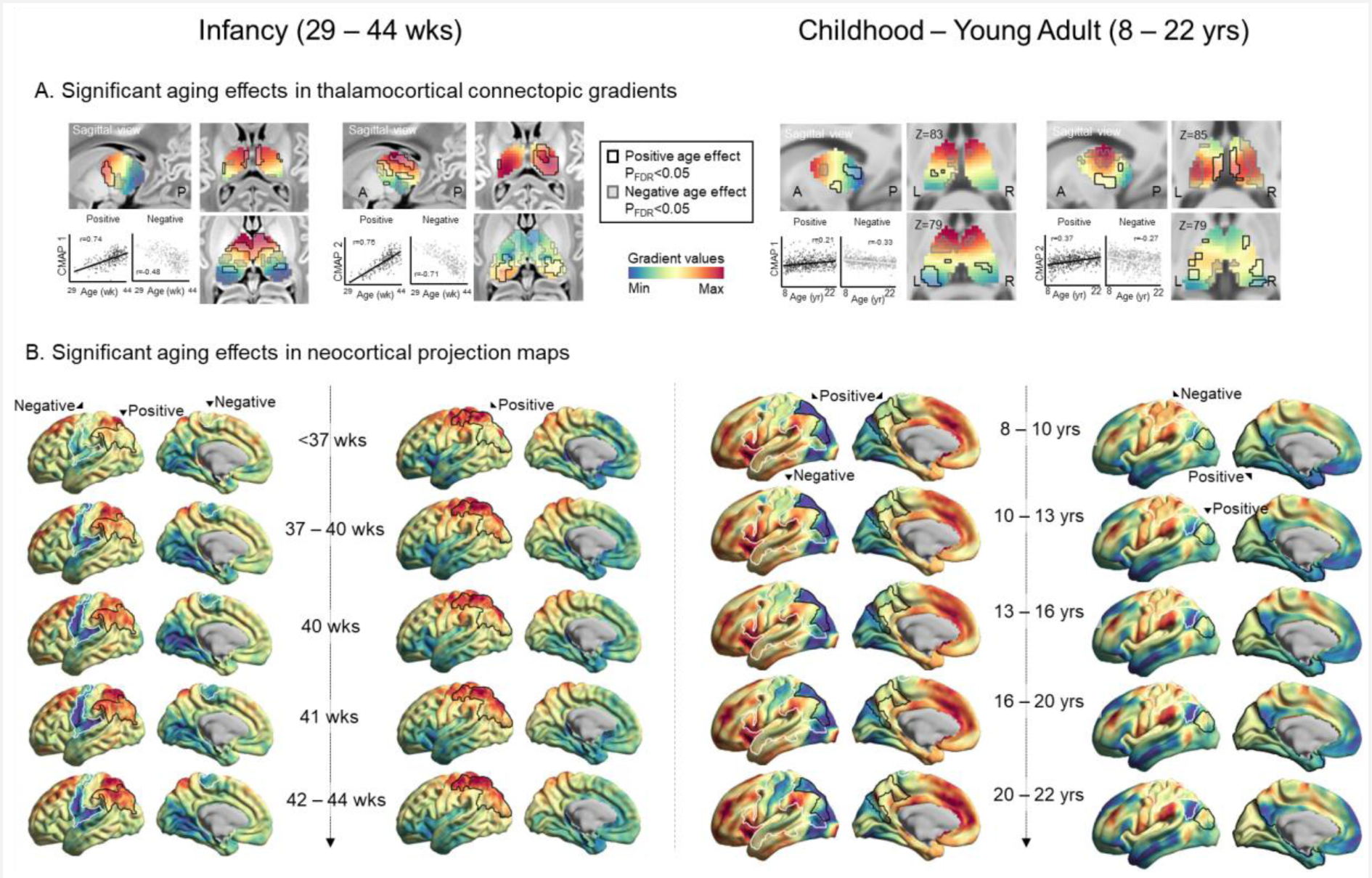
Age-related changes in (A) CMAPs and (B) NEOMAPs. Black lines delineate regions showing a significant positive aging effect while the white lines indicate a negative aging effect. Scatter plots illustrate the association between age and regions showing significant aging effects. Abbreviation, CMAP, thalamocortical connectopic map; NEOMAP, Neocortical projection map

**Supplementary Figure 5.**
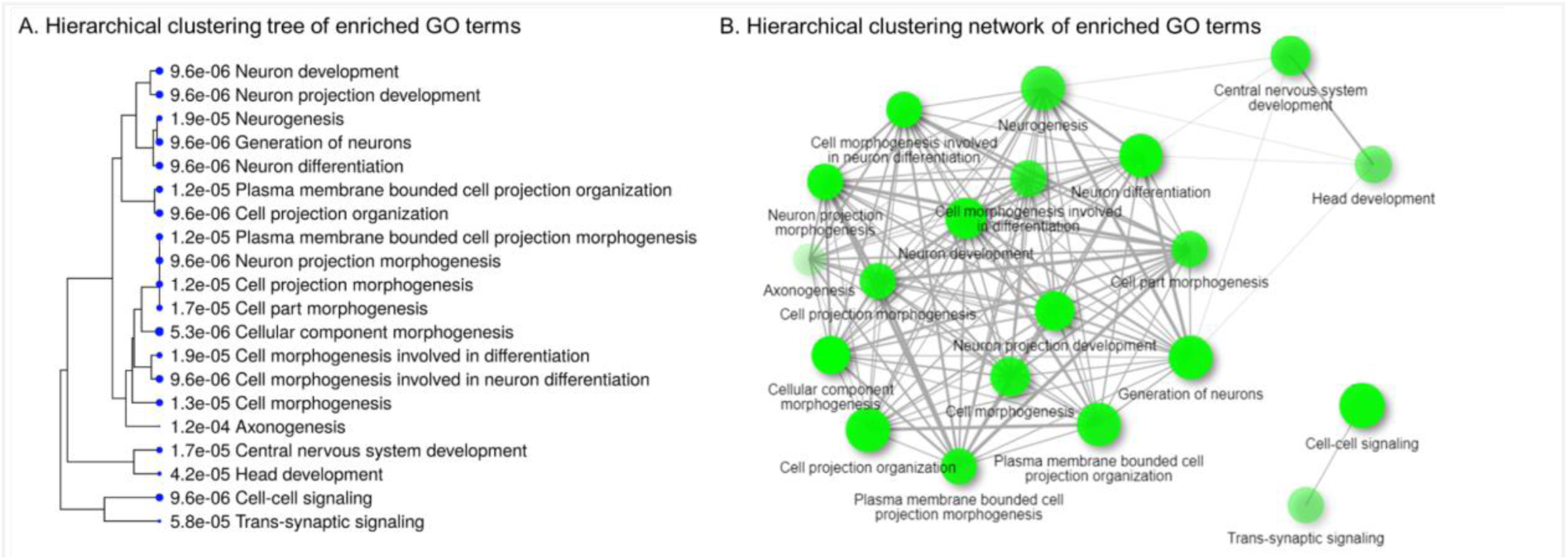
Relationships between the enriched GO biological process terms are shown as a (A) hierarchical clustering tree and (B) network. Pathways with many shared genes are clustered together. Two pathways (or nodes in the network) are connected if they share 20% or more genes. Darker nodes are more significantly enriched gene sets, while bigger nodes represent larger gene sets. Thicker edges represent more overlapped genes.

**Supplementary Figure 6.**
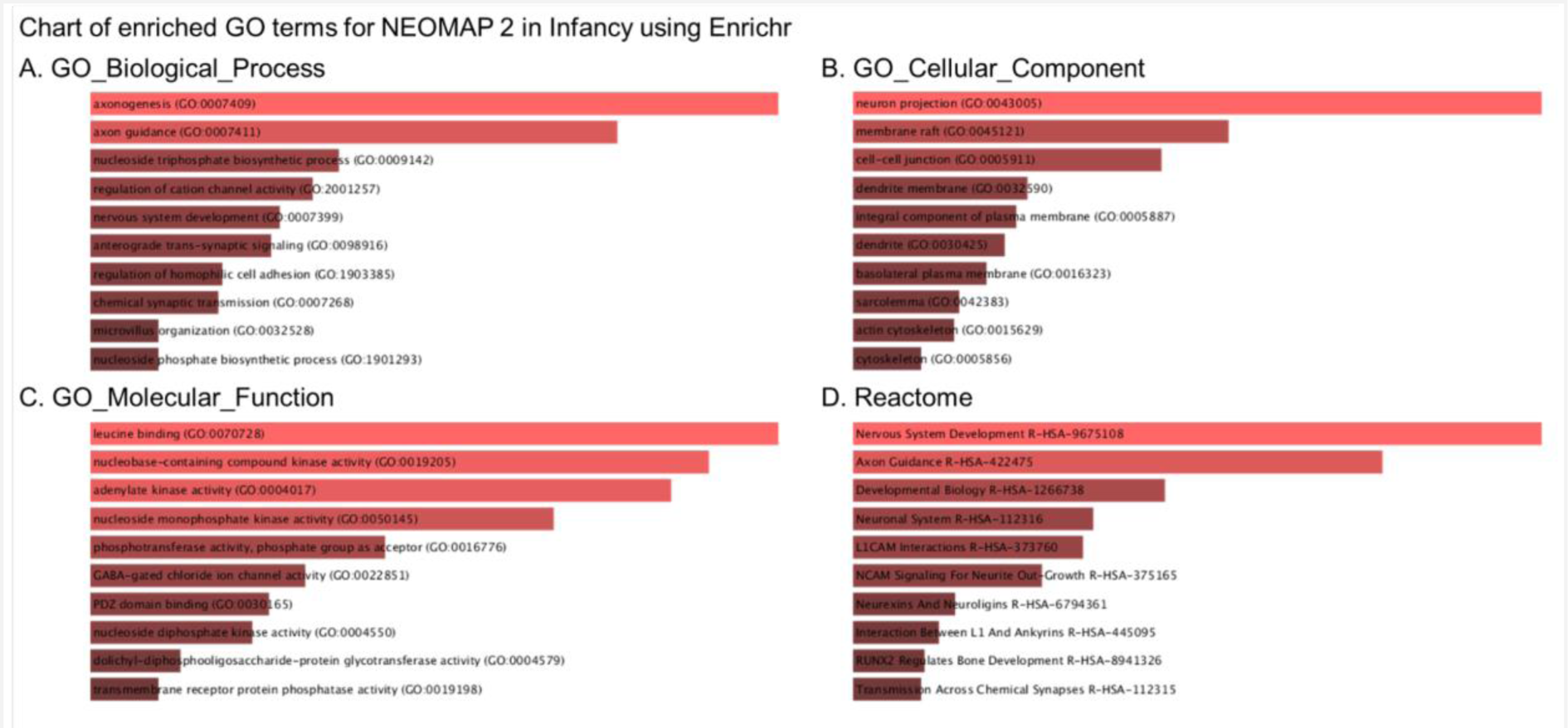
Gene ontology analysis using Enrichr largely replicated main results from ShinyGO. The gene ontology results for (A) biological processes, (B) cellular components, (C) molecular functions as well as data from the (D) reactome database are shown.

**Supplementary Figure 7.**
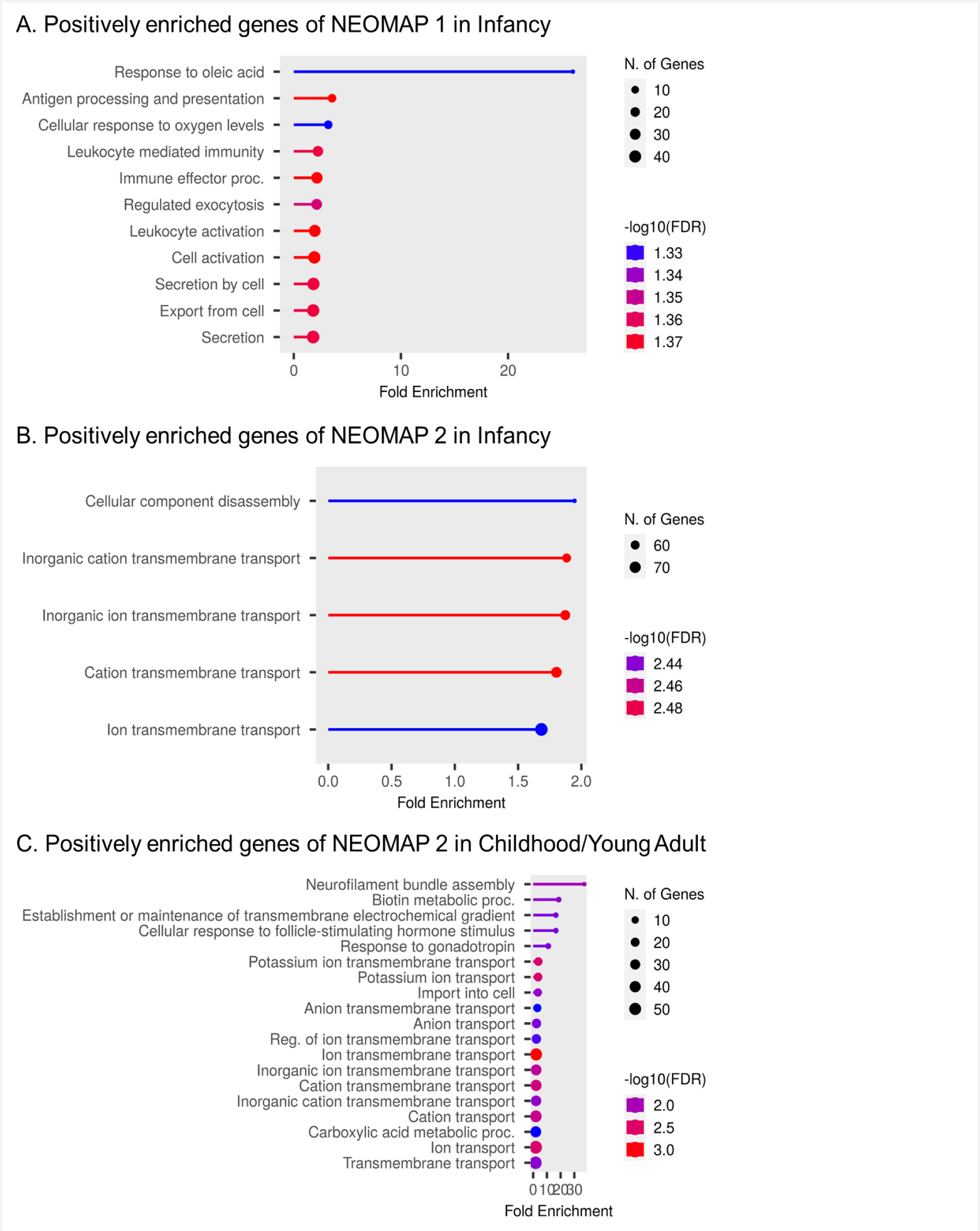
Chart of enriched gene ontology terms for NEOMAPs showing significant enrichment for (A) positively enriched genes of NEOMAP 1 in infancy, (B) positively enriched genes of NEOMAP2 in infancy, and (C) positively enriched genes of NEOMAP 2 in Childhood/Young adults.

**Supplementary Figure 8.**
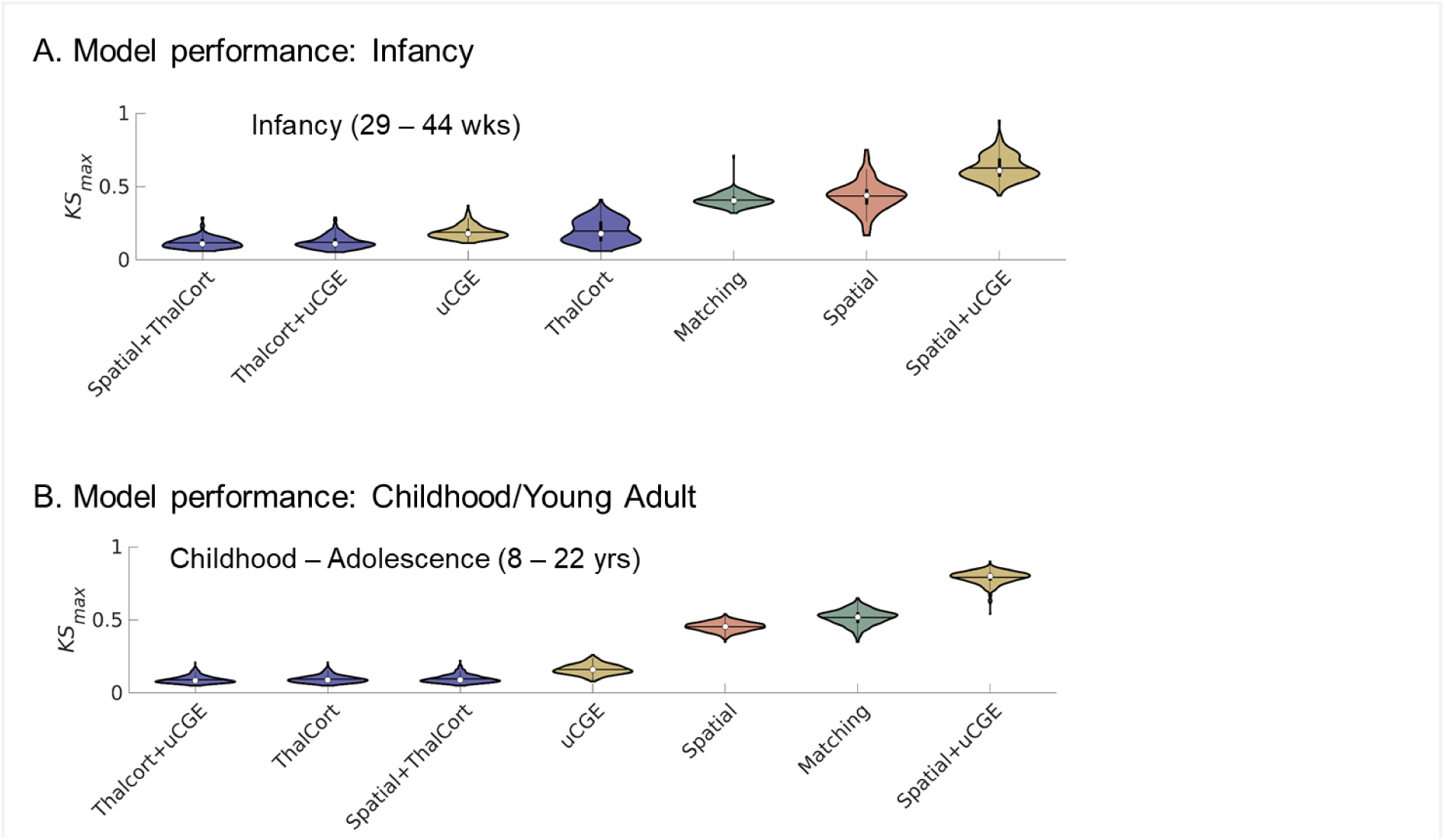
Model fitting performance for different wiring rules of generative network models for (A) infancy and (B) childhood/young adulthood.

**Supplementary Video.**
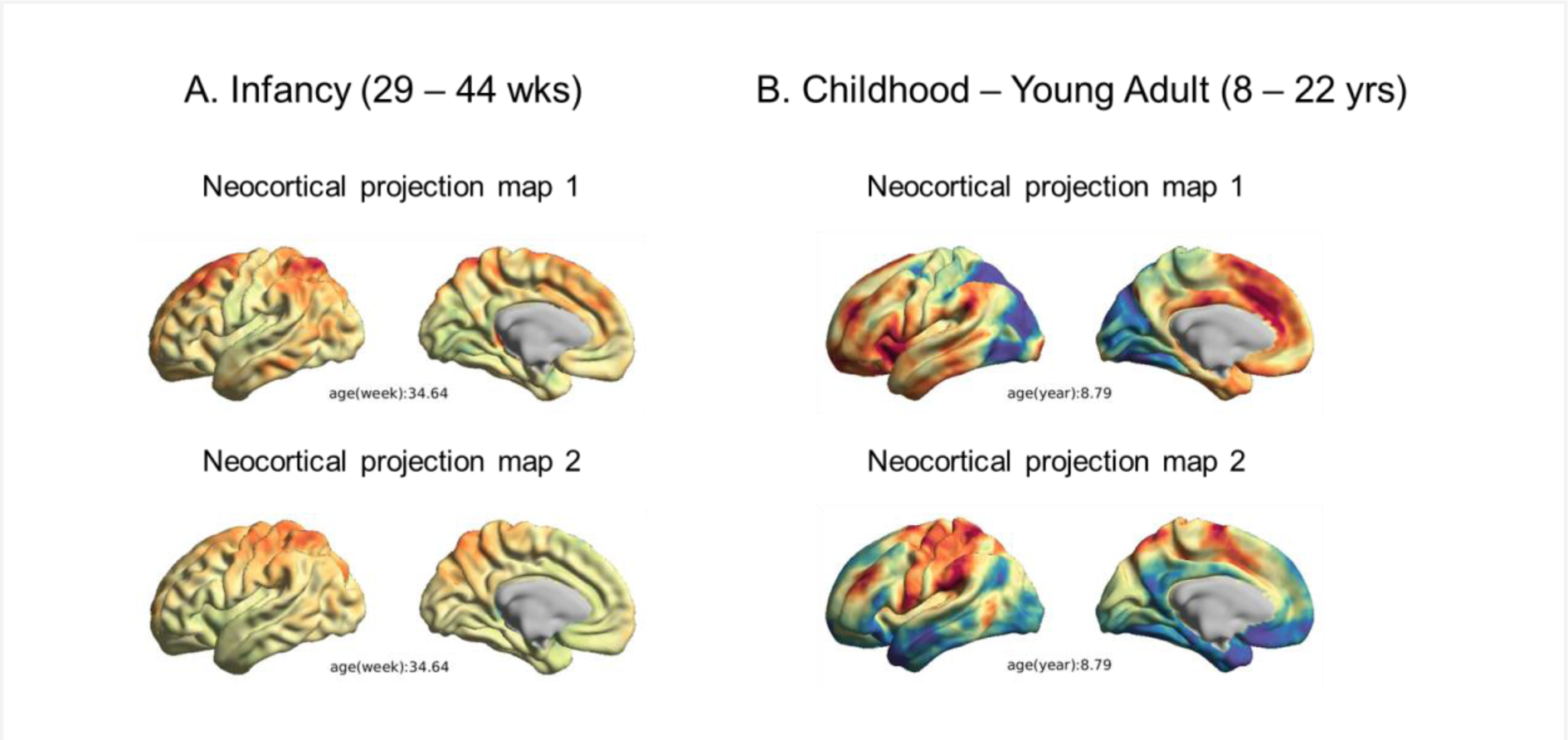

**Supplementary Table 1.** Biological processes of negative down-regulated genes from gene ontology enrichment analysis.

| Enrichment FDR | nGenes | Pathway Genes | Fold Enrichment | Pathways |
| --- | --- | --- | --- | --- |
| 9.6E-06 | 27 | 620 | 3.6 | Cell morphogenesis involved in neuron differentiation |
| 1.2E-04 | 21 | 490 | 3.5 | Axonogenesis |
| 9.6E-06 | 28 | 686 | 3.4 | Neuron projection morphogenesis |
| 5.3E-06 | 33 | 815 | 3.3 | Cellular component morphogenesis |
| 1.2E-05 | 28 | 700 | 3.3 | Plasma membrane bounded cell projection morphogenesis |
| 1.2E-05 | 28 | 704 | 3.3 | Cell projection morphogenesis |
| 1.7E-05 | 28 | 721 | 3.2 | Cell part morphogenesis |
| 1.9E-05 | 29 | 775 | 3.1 | Cell morphogenesis involved in differentiation |
| 5.8E-05 | 28 | 776 | 3 | Trans-synaptic signaling |
| 4.2E-05 | 30 | 856 | 2.9 | Head development |
| 9.6E-06 | 36 | 1045 | 2.8 | Neuron projection development |
| 1.3E-05 | 36 | 1085 | 2.7 | Cell morphogenesis |
| 1.7E-05 | 36 | 1105 | 2.7 | Central nervous system development |
| 9.6E-06 | 39 | 1203 | 2.7 | Neuron development |
| 9.6E-06 | 45 | 1473 | 2.5 | Neuron differentiation |
| 9.6E-06 | 49 | 1690 | 2.4 | Cell projection organization |
| 9.6E-06 | 47 | 1624 | 2.4 | Generation of neurons |
| 1.2E-05 | 47 | 1649 | 2.4 | Plasma membrane bounded cell projection organization |
| 9.6E-06 | 52 | 1885 | 2.3 | Cell-cell signaling |
| 1.9E-05 | 48 | 1757 | 2.3 | Neurogenesis |

**Supplementary Table 2.** Biological processes of positive up-regulated genes from gene ontology enrichment analysis.

| Enrichment FDR | nGenes | Pathway Genes | Fold Enrichment | Pathways |
| --- | --- | --- | --- | --- |
| 7.5E-03 | 5 | 17 | 12.9 | Neurotransmitter receptor transport to postsynaptic membrane |
| 7.5E-03 | 7 | 37 | 8.3 | Pos. reg. of sodium ion transport |
| 4.5E-03 | 8 | 45 | 7.8 | Mitochondrial fission |
| 7.5E-03 | 9 | 70 | 5.6 | Reg. of postsynaptic membrane neurotransmitter receptor levels |
| 5.4E-03 | 12 | 112 | 4.7 | Pos. reg. of ion transmembrane transporter activity |
| 2.9E-03 | 16 | 178 | 3.9 | Mitochondrial gene expression |
| 2.6E-03 | 20 | 261 | 3.4 | Protein complex oligomerization |
| 7.5E-03 | 17 | 229 | 3.3 | Insulin secretion |
| 7.5E-03 | 17 | 231 | 3.2 | Potassium ion transmembrane transport |
| 7.5E-03 | 18 | 255 | 3.1 | Potassium ion transport |
| 7.5E-03 | 19 | 274 | 3 | Peptide hormone secretion |
| 4.7E-04 | 37 | 615 | 2.6 | Cellular component disassembly |
| 3.4E-03 | 41 | 843 | 2.1 | Inorganic cation transmembrane transport |
| 2.6E-03 | 44 | 909 | 2.1 | Inorganic ion transmembrane transport |
| 2.6E-03 | 47 | 986 | 2.1 | Cation transmembrane transport |
| 2.6E-03 | 55 | 1251 | 1.9 | Ion transmembrane transport |
| 7.5E-03 | 67 | 1764 | 1.7 | Transmembrane transport |
| 7.5E-03 | 71 | 1910 | 1.6 | Reg. of transport |

## References

1. Gasser, R.F. (1975). Atlas of Human Embryos (HarperCollins Publishers).

2. Vogel, J.W., Alexander-Bloch, A., Wagstyl, K., Bertolero, M., Markello, R., Pines, A., Sydnor, V.J., Diaz-Papkovich, A., Hansen, J., Evans, A.C., et al. (2022). Conserved whole-brain spatiomolecular gradients shape adult brain functional organization. bioRxiv, 2022.09.18.508425. 10.1101/2022.09.18.508425.

3. O’Leary, D.D.M., and Nakagawa, Y. (2002). Patterning centers, regulatory genes and extrinsic mechanisms controlling arealization of the neocortex. Curr. Opin. Neurobiol. 12, 14–25.

4. Cadwell, C.R., Bhaduri, A., Mostajo-Radji, M.A., Keefe, M.G., and Nowakowski, T.J. (2019). Development and Arealization of the Cerebral Cortex. Neuron 103, 980–1004.

5. Rakic, P. (1988). Specification of cerebral cortical areas. Science 241, 170–176.

6. Rubenstein, J.L., and Rakic, P. (1999). Genetic control of cortical development. Cereb. Cortex 9, 521–523.

7. O’Leary, D.D. (1989). Do cortical areas emerge from a protocortex? Trends Neurosci. 12, 400–406.

8. Wolff, M., Morceau, S., Folkard, R., Martin-Cortecero, J., and Groh, A. (2021). A thalamic bridge from sensory perception to cognition. Neurosci. Biobehav. Rev. 120, 222–235.

9. Jones, E.G. (2012). The Thalamus (Springer Science & Business Media).

10. Müller, E.J., Munn, B., Hearne, L.J., Smith, J.B., Fulcher, B., Arnatkevičiūtė, A., Lurie, D.J., Cocchi, L., and Shine, J.M. (2020). Core and matrix thalamic sub-populations relate to spatio-temporal cortical connectivity gradients. Neuroimage 222, 117224.

11. Johnson, M.H. (2001). Functional brain development in humans. Nat. Rev. Neurosci. 2, 475–483.

12. Fox, S.E., Levitt, P., and Nelson, C.A., 3rd (2010). How the timing and quality of early experiences influence the development of brain architecture. Child Dev. 81, 28–40.

13. Tooley, U.A., Bassett, D.S., and Mackey, A.P. (2021). Environmental influences on the pace of brain development. Nat. Rev. Neurosci. 22, 372–384.

14. Nagalski, A., Puelles, L., Dabrowski, M., Wegierski, T., Kuznicki, J., and Wisniewska, M.B. (2016). Molecular anatomy of the thalamic complex and the underlying transcription factors. Brain Struct. Funct. 221, 2493–2510.

15. Gezelius, H., Moreno-Juan, V., Mezzera, C., Thakurela, S., Rodríguez-Malmierca, L.M., Pistolic, J., Benes, V., Tiwari, V.K., and López-Bendito, G. (2017). Genetic Labeling of Nuclei-Specific Thalamocortical Neurons Reveals Putative Sensory-Modality Specific Genes. Cereb. Cortex 27, 5054–5069.

16. Nakagawa, Y. (2019). Development of the thalamus: From early patterning to regulation of cortical functions. Wiley Interdiscip. Rev. Dev. Biol. 8, e345.

17. López-Bendito, G., and Molnár, Z. (2003). Thalamocortical development: how are we going to get there? Nat. Rev. Neurosci. 4, 276–289.

18. Molnár, Z., and Blakemore, C. (1995). How do thalamic axons find their way to the cortex? Trends Neurosci. 18, 389–397.

19. Sydnor, V.J., Larsen, B., Bassett, D.S., Alexander-Bloch, A., Fair, D.A., Liston, C., Mackey, A.P., Milham, M.P., Pines, A., Roalf, D.R., et al. (2021). Neurodevelopment of the association cortices: Patterns, mechanisms, and implications for psychopathology. Neuron 109, 2820–2846.

20. Tooley, U.A., Park, A.T., Leonard, J.A., Boroshok, A.L., McDermott, C.L., Tisdall, M.D., Bassett, D.S., and Mackey, A.P. (2022). The age of reason: Functional brain network development during childhood. J. Neurosci. 10.1523/JNEUROSCI.0511-22.2022.

21. Dong, H.-M., Margulies, D.S., Zuo, X.-N., and Holmes, A.J. (2021). Shifting gradients of macroscale cortical organization mark the transition from childhood to adolescence. Proc. Natl. Acad. Sci. U. S. A. 118. 10.1073/pnas.2024448118.

22. Park, B.-Y., Paquola, C., Bethlehem, R.A.I., Benkarim, O., Neuroscience in Psychiatry Network (NSPN) Consortium, Mišić, B., Smallwood, J., Bullmore, E.T., and Bernhardt, B.C. (2022). Adolescent development of multiscale structural wiring and functional interactions in the human connectome. Proc. Natl. Acad. Sci. U. S. A. 119, e2116673119.

23. Royer, J., Paquola, C., Larivière, S., Vos de Wael, R., Tavakol, S., Lowe, A.J., Benkarim, O., Evans, A.C., Bzdok, D., Smallwood, J., et al. (2020). Myeloarchitecture gradients in the human insula: Histological underpinnings and association to intrinsic functional connectivity. Neuroimage 216, 116859.

24. Alcauter, S., Lin, W., Smith, J.K., Short, S.J., Goldman, B.D., Reznick, J.S., Gilmore, J.H., and Gao, W. (2014). Development of thalamocortical connectivity during infancy and its cognitive correlations. J. Neurosci. 34, 9067–9075.

25. Fair, D.A., Bathula, D., Mills, K.L., Dias, T.G.C., Blythe, M.S., Zhang, D., Snyder, A.Z., Raichle, M.E., Stevens, A.A., Nigg, J.T., et al. (2010). Maturing thalamocortical functional connectivity across development. Front. Syst. Neurosci. 4, 10.

26. Larivière, S., Vos de Wael, R., Hong, S.-J., Paquola, C., Tavakol, S., Lowe, A.J., Schrader, D.V., and Bernhardt, B.C. (2020). Multiscale Structure–Function Gradients in the Neonatal Connectome. Cereb. Cortex 30, 47–58.

27. Haak, K.V., Marquand, A.F., and Beckmann, C.F. (2018). Connectopic mapping with resting-state fMRI. Neuroimage 170, 83–94.

28. Vos de Wael, R., Benkarim, O., Paquola, C., Lariviere, S., Royer, J., Tavakol, S., Xu, T., Hong, S.-J., Langs, G., Valk, S., et al. (2020). BrainSpace: a toolbox for the analysis of macroscale gradients in neuroimaging and connectomics datasets. Commun Biol 3, 103.

29. Margulies, D.S., Ghosh, S.S., Goulas, A., Falkiewicz, M., Huntenburg, J.M., Langs, G., Bezgin, G., Eickhoff, S.B., Castellanos, F.X., Petrides, M., et al. (2016). Situating the default-mode network along a principal gradient of macroscale cortical organization. Proc. Natl. Acad. Sci. U. S. A. 113, 12574–12579.

30. Van Essen, D.C., Smith, S.M., Barch, D.M., Behrens, T.E.J., Yacoub, E., Ugurbil, K., and WU-Minn HCP Consortium (2013). The WU-Minn Human Connectome Project: an overview. Neuroimage 80, 62–79.

31. Glasser, M.F., Sotiropoulos, S.N., Wilson, J.A., Coalson, T.S., Fischl, B., Andersson, J.L., Xu, J., Jbabdi, S., Webster, M., Polimeni, J.R., et al. (2013). The minimal preprocessing pipelines for the Human Connectome Project. Neuroimage 80, 105–124.

32. Makropoulos, A., Robinson, E.C., Schuh, A., Wright, R., Fitzgibbon, S., Bozek, J., Counsell, S.J., Steinweg, J., Vecchiato, K., Passerat-Palmbach, J., et al. (2018). The developing human connectome project: A minimal processing pipeline for neonatal cortical surface reconstruction. Neuroimage 173, 88–112.

33. Fitzgibbon, S.P., Harrison, S.J., Jenkinson, M., Baxter, L., Robinson, E.C., Bastiani, M., Bozek, J., Karolis, V., Cordero Grande, L., Price, A.N., et al. (2020). The developing Human Connectome Project (dHCP) automated resting-state functional processing framework for newborn infants. Neuroimage 223, 117303.

34. Somerville, L.H., Bookheimer, S.Y., Buckner, R.L., Burgess, G.C., Curtiss, S.W., Dapretto, M., Elam, J.S., Gaffrey, M.S., Harms, M.P., Hodge, C., et al. (2018). The Lifespan Human Connectome Project in Development: A large-scale study of brain connectivity development in 5–21 year olds. Neuroimage 183, 456–468.

35. Yeo, B.T.T., Krienen, F.M., Sepulcre, J., Sabuncu, M.R., Lashkari, D., Hollinshead, M., Roffman, J.L., Smoller, J.W., Zöllei, L., Polimeni, J.R., et al. (2011). The organization of the human cerebral cortex estimated by intrinsic functional connectivity. J. Neurophysiol. 106, 1125–1165.

36. Hawrylycz, M.J., Lein, E.S., Guillozet-Bongaarts, A.L., Shen, E.H., Ng, L., Miller, J.A., van de Lagemaat, L.N., Smith, K.A., Ebbert, A., Riley, Z.L., et al. (2012). An anatomically comprehensive atlas of the adult human brain transcriptome. Nature 489, 391–399.

37. Raut, R.V., Snyder, A.Z., and Raichle, M.E. (2020). Hierarchical dynamics as a macroscopic organizing principle of the human brain. Proc. Natl. Acad. Sci. U. S. A. 117, 20890–20897.

38. Menon, V., and Uddin, L.Q. (2010). Saliency, switching, attention and control: a network model of insula function. Brain Struct. Funct. 214, 655–667.

39. Sepulcre, J., Liu, H., Talukdar, T., Martincorena, I., Yeo, B.T.T., and Buckner, R.L. (2010). The organization of local and distant functional connectivity in the human brain. PLoS Comput. Biol. 6, e1000808.

40. Jones, E.G. (1998). Viewpoint: the core and matrix of thalamic organization. Neuroscience 85, 331–345.

41. Ge, S.X., Jung, D., and Yao, R. (2020). ShinyGO: a graphical gene-set enrichment tool for animals and plants. Bioinformatics 36, 2628–2629.

42. Dougherty, J.D., Schmidt, E.F., Nakajima, M., and Heintz, N. (2010). Analytical approaches to RNA profiling data for the identification of genes enriched in specific cells. Nucleic Acids Res. 38, 4218–4230.

43. Kuleshov, M.V., Jones, M.R., Rouillard, A.D., Fernandez, N.F., Duan, Q., Wang, Z., Koplev, S., Jenkins, S.L., Jagodnik, K.M., Lachmann, A., et al. (2016). Enrichr: a comprehensive gene set enrichment analysis web server 2016 update. Nucleic Acids Res. 44, W90–W97.

44. Chen, E.Y., Tan, C.M., Kou, Y., Duan, Q., Wang, Z., Meirelles, G.V., Clark, N.R., and Ma’ayan, A. (2013). Enrichr: interactive and collaborative HTML5 gene list enrichment analysis tool. BMC Bioinformatics 14, 128.

45. Xie, Z., Bailey, A., Kuleshov, M.V., Clarke, D.J.B., Evangelista, J.E., Jenkins, S.L., Lachmann, A., Wojciechowicz, M.L., Kropiwnicki, E., Jagodnik, K.M., et al. (2021). Gene Set Knowledge Discovery with Enrichr. Curr Protoc 1, e90.

46. Betzel, R.F., and Bassett, D.S. (2017). Generative models for network neuroscience: prospects and promise. J. R. Soc. Interface 14. 10.1098/rsif.2017.0623.

47. Oldham, S., Fulcher, B.D., Aquino, K., Arnatkevičiūtė, A., Paquola, C., Shishegar, R., and Fornito, A. (2022). Modeling spatial, developmental, physiological, and topological constraints on human brain connectivity. Sci Adv 8, eabm6127.

48. Betzel, R.F., Avena-Koenigsberger, A., Goñi, J., He, Y., de Reus, M.A., Griffa, A., Vértes, P.E., Mišic, B., Thiran, J.-P., Hagmann, P., et al. (2016). Generative models of the human connectome. Neuroimage 124, 1054–1064.

49. Paus, T. (2005). Mapping brain maturation and cognitive development during adolescence. Trends Cogn. Sci. 9, 60–68.

50. Hwang, K., Bertolero, M.A., Liu, W.B., and D’Esposito, M. (2017). The Human Thalamus Is an Integrative Hub for Functional Brain Networks. J. Neurosci. 37, 5594–5607.

51. Ball, G., Srinivasan, L., Aljabar, P., Counsell, S.J., Durighel, G., Hajnal, J.V., Rutherford, M.A., and Edwards, A.D. (2013). Development of cortical microstructure in the preterm human brain. Proc. Natl. Acad. Sci. U. S. A. 110, 9541–9546.

52. Baum, G.L., Cui, Z., Roalf, D.R., and Ciric, R. (2020). Development of structure–function coupling in human brain networks during youth. Proceedings of the.

53. Mitchell, A.S. (2015). The mediodorsal thalamus as a higher order thalamic relay nucleus important for learning and decision-making. Neurosci. Biobehav. Rev. 54, 76– 88.

54. Toulmin, H., O’Muircheartaigh, J., Counsell, S.J., Falconer, S., Chew, A., Beckmann, C.F., and Edwards, A.D. (2021). Functional thalamocortical connectivity at term equivalent age and outcome at 2 years in infants born preterm. Cortex 135, 17–29.

55. Huttenlocher, P.R., and Dabholkar, A.S. (1997). Regional differences in synaptogenesis in human cerebral cortex. J. Comp. Neurol. 387, 167–178.

56. Dosenbach, N.U.F., Nardos, B., Cohen, A.L., Fair, D.A., Power, J.D., Church, J.A., Nelson, S.M., Wig, G.S., Vogel, A.C., Lessov-Schlaggar, C.N., et al. (2010). Prediction of individual brain maturity using fMRI. Science 329, 1358–1361.

57. Petanjek, Z., Judas, M., Kostović, I., and Uylings, H.B.M. (2008). Lifespan alterations of basal dendritic trees of pyramidal neurons in the human prefrontal cortex: a layer-specific pattern. Cereb. Cortex 18, 915–929.

58. Valk, S.L., Xu, T., Paquola, C., Park, B.-Y., Bethlehem, R.A.I., Vos de Wael, R., Royer, J., Masouleh, S.K., Bayrak, Ş., Kochunov, P., et al. (2022). Genetic and phylogenetic uncoupling of structure and function in human transmodal cortex. Nat. Commun. 13, 2341.

59. Geschwind, D.H., and Rakic, P. (2013). Cortical evolution: judge the brain by its cover. Neuron 80, 633–647.

60. Buckner, R.L., and Krienen, F.M. (2013). The evolution of distributed association networks in the human brain. Trends Cogn. Sci. 17, 648–665.

61. Blumenfeld, H. (2016). Chapter 1 - Neuroanatomical Basis of Consciousness. In The Neurology of Conciousness (Second Edition), S. Laureys, O. Gosseries, and G. Tononi, eds. (Academic Press), pp. 3–29.

62. Innocenti, G.M. (1995). Exuberant development of connections, and its possible permissive role in cortical evolution. Trends Neurosci. 18, 397–402.

63. Bourgeois, J.P. (1997). Synaptogenesis, heterochrony and epigenesis in the mammalian neocortex. Acta Paediatr. Suppl. 422, 27–33.

64. Hill, J., Inder, T., Neil, J., Dierker, D., Harwell, J., and Van Essen, D. (2010). Similar patterns of cortical expansion during human development and evolution. Proc. Natl. Acad. Sci. U. S. A. 107, 13135–13140.

65. Grayson, D.S., and Fair, D.A. (2017). Development of large-scale functional networks from birth to adulthood: A guide to the neuroimaging literature. Neuroimage 160, 15–31.

66. Petanjek, Z., Judaš, M., Šimic, G., Rasin, M.R., Uylings, H.B.M., Rakic, P., and Kostovic, I. (2011). Extraordinary neoteny of synaptic spines in the human prefrontal cortex. Proc. Natl. Acad. Sci. U. S. A. 108, 13281–13286.

67. Stiles, J., and Jernigan, T.L. (2010). The basics of brain development. Neuropsychol. Rev. 20, 327–348.

68. Huttenlocher, P.R. (1979). Synaptic density in human frontal cortex - developmental changes and effects of aging. Brain Res. 163, 195–205.

69. Kast, R.J., and Levitt, P. (2019). Precision in the development of neocortical architecture: From progenitors to cortical networks. Prog. Neurobiol. 175, 77–95.

70. Vue, T.Y., Lee, M., Tan, Y.E., Werkhoven, Z., Wang, L., and Nakagawa, Y. (2013). Thalamic control of neocortical area formation in mice. J. Neurosci. 33, 8442–8453.

71. Goulas, A., Margulies, D.S., Bezgin, G., and Hilgetag, C.C. (2019). The architecture of mammalian cortical connectomes in light of the theory of the dual origin of the cerebral cortex. Cortex 118, 244–261.

72. Valk, S.L., Xu, T., Margulies, D.S., Masouleh, S.K., Paquola, C., Goulas, A., Kochunov, P., Smallwood, J., Yeo, B.T.T., Bernhardt, B.C., et al. (2020). Shaping brain structure: Genetic and phylogenetic axes of macroscale organization of cortical thickness. Sci Adv 6. 10.1126/sciadv.abb3417.

73. Pandya, D., Petrides, M., and Cipolloni, P.B. (2015). Cerebral Cortex: Architecture, Connections, and the Dual Origin Concept (Oxford University Press).

74. Hilgetag, C.C., Goulas, A., and Changeux, J.-P. (2022). A natural cortical axis connecting the outside and inside of the human brain. Netw Neurosci 6, 950–959.

75. Corbetta, M., and Shulman, G.L. (2002). Control of goal-directed and stimulus-driven attention in the brain. Nat. Rev. Neurosci. 3, 201–215.

76. Fox, M.D., Corbetta, M., Snyder, A.Z., Vincent, J.L., and Raichle, M.E. (2006). Spontaneous neuronal activity distinguishes human dorsal and ventral attention systems. Proc. Natl. Acad. Sci. U. S. A. 103, 10046–10051.

77. Vossel, S., Geng, J.J., and Fink, G.R. (2014). Dorsal and ventral attention systems: distinct neural circuits but collaborative roles. Neuroscientist 20, 150–159.

78. Corcoran, K.A., Frick, B.J., Radulovic, J., and Kay, L.M. (2016). Analysis of coherent activity between retrosplenial cortex, hippocampus, thalamus, and anterior cingulate cortex during retrieval of recent and remote context fear memory. Neurobiol. Learn. Mem. 127, 93–101.

79. Halassa, M.M., and Kastner, S. (2017). Thalamic functions in distributed cognitive control. Nat. Neurosci. 20, 1669–1679.

80. Wolff, M., and Vann, S.D. (2019). The Cognitive Thalamus as a Gateway to Mental Representations. J. Neurosci. 39, 3–14.

81. Keller, A.S., Sydnor, V.J., Pines, A., Fair, D.A., Bassett, D.S., and Satterthwaite, T.D. (2023). Hierarchical functional system development supports executive function. Trends Cogn. Sci. 27, 160–174.

82. Seeley, W.W. (2019). The Salience Network: A Neural System for Perceiving and Responding to Homeostatic Demands. J. Neurosci. 39, 9878–9882.

83. Vértes, P.E., Alexander-Bloch, A.F., Gogtay, N., Giedd, J.N., Rapoport, J.L., and Bullmore, E.T. (2012). Simple models of human brain functional networks. Proc. Natl. Acad. Sci. U. S. A. 109, 5868–5873.

84. Liu, X., and Duyn, J.H. (2013). Time-varying functional network information extracted from brief instances of spontaneous brain activity. Proc. Natl. Acad. Sci. U. S. A. 110, 4392–4397.

85. Majeed, W., Magnuson, M., Hasenkamp, W., Schwarb, H., Schumacher, E.H., Barsalou, L., and Keilholz, S.D. (2011). Spatiotemporal dynamics of low frequency BOLD fluctuations in rats and humans. Neuroimage 54, 1140–1150.

86. Bell, P.T., and Shine, J.M. (2016). Subcortical contributions to large-scale network communication. Neurosci. Biobehav. Rev. 71, 313–322.

87. Shine, J.M. (2021). The thalamus integrates the macrosystems of the brain to facilitate complex, adaptive brain network dynamics. Prog. Neurobiol. 199, 101951.

88. Hintzen, A., Pelzer, E.A., and Tittgemeyer, M. (2018). Thalamic interactions of cerebellum and basal ganglia. Brain Struct. Funct.

89. Caciagli, L., Allen, L.A., He, X., Trimmel, K., Vos, S.B., Centeno, M., Galovic, M., Sidhu, M.K., Thompson, P.J., Bassett, D.S., et al. (2020). Thalamus and focal to bilateral seizures: A multiscale cognitive imaging study. Neurology 95, e2427–e2441.

90. Weng, Y., Larivière, S., Caciagli, L., Vos de Wael, R., Rodríguez-Cruces, R., Royer, J., Xu, Q., Bernasconi, N., Bernasconi, A., Thomas Yeo, B.T., et al. (2020). Macroscale and microcircuit dissociation of focal and generalized human epilepsies. Commun Biol 3, 244.

91. Bernhardt, B.C., Bernasconi, N., Kim, H., and Bernasconi, A. (2012). Mapping thalamocortical network pathology in temporal lobe epilepsy. Neurology 78, 129–136.

92. Hwang, W.J., Kwak, Y.B., Cho, K.I.K., Lee, T.Y., Oh, H., Ha, M., Kim, M., and Kwon, J.S. (2022). Thalamic Connectivity System Across Psychiatric Disorders: Current Status and Clinical Implications. Biol Psychiatry Glob Open Sci 2, 332–340.

93. Park, B.-Y., Hong, S.-J., Valk, S.L., Paquola, C., Benkarim, O., Bethlehem, R.A.I., Di Martino, A., Milham, M.P., Gozzi, A., Yeo, B.T.T., et al. (2021). Differences in subcortico-cortical interactions identified from connectome and microcircuit models in autism. Nat. Commun. 12, 2225.

94. Park, S., Haak, K.V., Cho, H.B., Valk, S.L., Bethlehem, R.A.I., Milham, M.P., Bernhardt, B.C., Di Martino, A., and Hong, S.-J. (2021). Atypical Integration of Sensory-to-Transmodal Functional Systems Mediates Symptom Severity in Autism. Front. Psychiatry 12, 699813.

95. Smith, S.M. (2002). Fast robust automated brain extraction. Hum. Brain Mapp. 17, 143– 155.

96. Makropoulos, A., Gousias, I.S., Ledig, C., Aljabar, P., Serag, A., Hajnal, J.V., Edwards, A.D., Counsell, S.J., and Rueckert, D. (2014). Automatic whole brain MRI segmentation of the developing neonatal brain. IEEE Trans. Med. Imaging 33, 1818–1831.

97. Robinson, E.C., Jbabdi, S., Glasser, M.F., Andersson, J., Burgess, G.C., Harms, M.P., Smith, S.M., Van Essen, D.C., and Jenkinson, M. (2014). MSM: a new flexible framework for Multimodal Surface Matching. Neuroimage 100, 414–426.

98. Robinson, E.C., Jbabdi, S., Andersson, J., Smith, S., Glasser, M.F., Van Essen, D.C., Burgess, G., Harms, M.P., Barch, D.M., and Barch, D.M. (2013). Multimodal surface matching: fast and generalisable cortical registration using discrete optimisation. Inf. Process. Med. Imaging 23, 475–486.

99. Tian, Y., Margulies, D.S., Breakspear, M., and Zalesky, A. (2020). Topographic organization of the human subcortex unveiled with functional connectivity gradients. Nat. Neurosci. 23, 1421–1432.

100. Marquand, A.F., Haak, K.V., and Beckmann, C.F. (2017). Functional corticostriatal connection topographies predict goal directed behaviour in humans. Nat Hum Behav 1, 0146.

101. Larivière, S., Bayrak, Ş., Vos de Wael, R., Benkarim, O., Herholz, P., Rodriguez-Cruces, R., Paquola, C., Hong, S.-J., Misic, B., Evans, A.C., et al. (2023). BrainStat: A toolbox for brain-wide statistics and multimodal feature associations. Neuroimage 266, 119807.

102. Worsley, K.J., Taylor, J., Carbonell, F., Chung, M., Duerden, E., Bernhardt, B., Lyttelton, O., Boucher, M., and Evans, A. (2009). A Matlab toolbox for the statistical analysis of univariate and multivariate surface and volumetric data using linear mixed effects models and random field theory. In NeuroImage Organisation for Human Brain Mapping 2009 Annual Meeting (math.mcgill.ca), p. S102.

103. Kilford, E.J., Garrett, E., and Blakemore, S.-J. (2016). The development of social cognition in adolescence: An integrated perspective. Neurosci. Biobehav. Rev. 70, 106– 120.

104. Güroğlu, B., van den Bos, W., and Crone, E.A. (2014). Sharing and giving across adolescence: an experimental study examining the development of prosocial behavior. Front. Psychol. 5, 291.

105. Keshavan, M.S., Diwadkar, V.A., DeBellis, M., Dick, E., Kotwal, R., Rosenberg, D.R., Sweeney, J.A., Minshew, N., and Pettegrew, J.W. (2002). Development of the corpus callosum in childhood, adolescence and early adulthood. Life Sci. 70, 1909–1922.

106. Arnatkeviciute, A., Fulcher, B.D., and Fornito, A. (2019). A practical guide to linking brain-wide gene expression and neuroimaging data. Neuroimage 189, 353–367.

107. Quackenbush, J. (2002). Microarray data normalization and transformation. Nat. Genet. 32 *Suppl*, 496–501.

108. Hawrylycz, M., Miller, J.A., Menon, V., Feng, D., Dolbeare, T., Guillozet-Bongaarts, A.L., Jegga, A.G., Aronow, B.J., Lee, C.-K., Bernard, A., et al. (2015). Canonical genetic signatures of the adult human brain. Nat. Neurosci. 18, 1832–1844.

109. Fulcher, B.D., Little, M.A., and Jones, N.S. (2013). Highly comparative time-series analysis: the empirical structure of time series and their methods. J. R. Soc. Interface 10, 20130048.

110. Burt, J.B., Helmer, M., Shinn, M., Anticevic, A., and Murray, J.D. (2020). Generative modeling of brain maps with spatial autocorrelation. Neuroimage 220, 117038.

111. Fontana, L., Partridge, L., and Longo, V.D. (2010). Extending Healthy Life Span—From Yeast to Humans. Science 328, 321–326.

112. Parikh, A., Miranda, E.R., Katoh-Kurasawa, M., Fuller, D., Rot, G., Zagar, L., Curk, T., Sucgang, R., Chen, R., Zupan, B., et al. (2010). Conserved developmental transcriptomes in evolutionarily divergent species. Genome Biol. 11, R35.

113. Arnatkeviciute, A., Fulcher, B.D., Oldham, S., Tiego, J., Paquola, C., Gerring, Z., Aquino, K., Hawi, Z., Johnson, B., Ball, G., et al. (2021). Genetic influences on hub connectivity of the human connectome. Nat. Commun. 12, 4237.

114. Akarca, D., Vértes, P.E., Bullmore, E.T., CALM team, and Astle, D.E. (2021). A generative network model of neurodevelopmental diversity in structural brain organization. Nat. Commun. 12, 4216.

115. Zhang, X., Braun, U., Harneit, A., Zang, Z., Geiger, L.S., Betzel, R.F., Chen, J., Schweiger, J., Schwarz, K., Reinwald, J.R., et al. Generative network models identify biological mechanisms of altered structural brain connectivity in schizophrenia. 10.1101/604322.

116. Yang, S., Wagstyl, K., Meng, Y., Zhao, X., Li, J., Zhong, P., Li, B., Fan, Y.-S., Chen, H., and Liao, W. (2021). Cortical patterning of morphometric similarity gradient reveals diverged hierarchical organization in sensory-motor cortices. Cell Rep. 36, 109582.

117. Burt, J.B., Demirtaş, M., Eckner, W.J., Navejar, N.M., Ji, J.L., Martin, W.J., Bernacchia, A., Anticevic, A., and Murray, J.D. (2018). Hierarchy of transcriptomic specialization across human cortex captured by structural neuroimaging topography. Nat. Neurosci. 21, 1251–1259.

118. Fornito, A., Zalesky, A., and Bullmore, E. (2016). Fundamentals of Brain Network Analysis (Academic Press).

119. Rubinov, M., and Sporns, O. (2010). Complex network measures of brain connectivity: uses and interpretations. Neuroimage 52, 1059–1069.

120. Liu, Y., Seguin, C., Mansour, S., Oldham, S., Betzel, R., Biase, M.D., and Zalesky, A. (2023). Parameter estimation for connectome generative models: Accuracy, reliability, and a fast parameter fitting method. Neuroimage, 119962.

121. Mitra, A., Snyder, A.Z., Tagliazucchi, E., Laufs, H., Elison, J., Emerson, R.W., Shen, M.D., Wolff, J.J., Botteron, K.N., Dager, S., et al. (2017). Resting-state fMRI in sleeping infants more closely resembles adult sleep than adult wakefulness. PLoS One 12, e0188122.

122. Larson-Prior, L.J., Power, J.D., Vincent, J.L., Nolan, T.S., Coalson, R.S., Zempel, J., Snyder, A.Z., Schlaggar, B.L., Raichle, M.E., and Petersen, S.E. (2011). Modulation of the brain’s functional network architecture in the transition from wake to sleep. Slow Brain Oscillations of Sleep, Resting State and Vigilance, 277–294. 10.1016/b978-0-444-53839-0.00018-1.

123. Spoormaker, V.I., Gleiser, P.M., and Czisch, M. (2012). Frontoparietal Connectivity and Hierarchical Structure of the Brain’s Functional Network during Sleep. Frontiers in Neurology 3. 10.3389/fneur.2012.00080.

124. Horovitz, S.G., Fukunaga, M., de Zwart, J.A., van Gelderen, P., Fulton, S.C., Balkin, T.J., and Duyn, J.H. (2008). Low frequency BOLD fluctuations during resting wakefulness and light sleep: a simultaneous EEG-fMRI study. Hum. Brain Mapp. 29, 671–682.

125. Gao, W., Gilmore, J.H., Shen, D., Smith, J.K., Zhu, H., and Lin, W. (2013). The synchronization within and interaction between the default and dorsal attention networks in early infancy. Cereb. Cortex 23, 594–603.

126. Sharad, S., Brian, C., Ranjit, K., Satra, G., Chao-gan, Y., Qingyang, L., Joshua, V., Randal, B., Stanley, C., Cameron, C., et al. (2014). Towards Automated Analysis of Connectomes: The Configurable Pipeline for the Analysis of Connectomes (C-PAC). Frontiers in Neuroinformatics 8. 10.3389/conf.fninf.2014.08.00117.

